# Life in a Central European warm-temperate to subtropical open forest: paleoecology of the rhinocerotids from Ulm-Westtangente (Aquitanian, early Miocene, Germany)

**DOI:** 10.1101/2023.09.22.558985

**Authors:** Manon Hullot, Céline Martin, Cécile Blondel, Gertrud Rößner

**Affiliations:** Bayerische Staatssammlung für Paläontologie und Geologie, Richard-Wagner Straße, 10 8033 München, Germany; Géosciences Montpellier, Université de Montpellier Bât 22 – Place Eugène Bataillon, 34090, Montpellier, France; PALEVOPRIM Poitiers, Université de Poitiers Bât B35 – TSA 51106, 6 rue Michel Brunet, 86073, Poitiers, France; Department für Geo- und Umweltwissenschaften, Paläontologie & Geobiologie, Ludwig-Maximilians-Universität München, Richard-Wagner-Strasse 10, 80333, Munich, Germany

**Keywords:** diet, habitat, niche partioning, Freshwater Molasse Germany

## Abstract

The locality of Ulm-Westtangente yielded the richest vertebrate fauna from the Aquitanian of Germany. Its dating to the Mammal Neogene Zone 2a, a turnover in Cenozoic climate, makes it a crucial source for the understanding of faunal, palaeoecological and palaeoenvironmental specifics of the European Aquitanian. However, if most taxa from Ulm-Westtangente have been studied, very little to nothing has been done on the large herbivores and notably on the two rhinocerotids *Mesaceratherium paulhiacense* and *Protaceratherium minutum*. Here, we used a multi-proxy approach to investigate the paleoecology of these two species. The remains of the smaller species *P. minutum* (442 to 667 kg) are twice as abundant as those of the larger *M. paulhiacense* (1687 to 2576 kg), but both display a similar age structure (∼ 10 % of juveniles, 20 % of subadults and 70 % of adults), mortality curves, and mild prevalence of hypoplasia (∼ 17 %). Results from dental mesowear, microwear, and carbon isotopes indicate different feeding preferences: both were C3 feeders but *M. paulhiacense* had a more abrasive diet and was probably a mixed feeder. Our study on rhinocerotids also yielded new paleoenvironmental insights, such as the mean annual temperature (15.8 °C) and precipitation (317 mm/year) suggesting rather warm and dry conditions.

**Statements and Declarations:** The authors have no relevant financial or non-financial interests to disclose. This study was funded by a post doctoral fellowship of the Alexander von Humboldt Foundation (Germany).

## Introduction

The early Miocene is a key period in Rhinocerotidae evolution, as it was a time of great diversification and geographical expansion. Indeed, the family experienced a peak of alpha-diversity during late early Miocene with several sympatric species at single fossil sites (Antoine and Becker 2013). The early Miocene is also marked by a turnover and a great endemism in rhinoceros species of Western Europe. Despite all these, the paleoecology of rhinocerotids during this important time period has rarely been studied, although it has the potential to unravel niche partioning and ecological shifts associated with climatic conditions (Hullot et al. 2023a, b).

One of the richest fossil mammal localities of the early Miocene (Aquitanian: 23.04 – 20.44 Mya; Raffi et al. 2020) in Europe is Ulm-Westtangente in Germany (Heizmann et al. 1989; Costeur et al. 2012). Ulm-Westtangente is located in the Lower Freshwater Molasse sediments of the Baden-Württemberg Basin in Southwestern Germany, about 5 km North-West of Ulm (590 m above sea level, coordinates: 48.418321 N, 9.933701 E – converted from the Gauss Kruger coordinates in original publication of Heizmann et al. 1989: r 35 69 188, h 53 64 925). It has been dated to the early Miocene by correlation with the Mammal Neogene-Zone 2a (MN2a; Bruijn et al. 1992; Steininger 1999), an interval with relatively warm and stable climatic conditions following the cold start of the Miocene (MN1) due to the Mi-1 glaciation event (Zachos et al. 2001; Westerhold et al. 2020). A single 35 cm-thick layer has yielded more than 60 mammal species, making it the richest vertebrate fauna from the early Miocene ever found in Germany (Heizmann et al. 1989; Costeur et al. 2012). Among the numerous fossil remains excavated (∼ 6000 from large mammals, > 6000 from small mammals and > 1000 from other vertebrates; Heizmann et al. 1989), abundant material has been attributed to two species of rhinocerotids: the small tapir-sized *Protaceratherium minutum* and the medium to large sized *Mesaceratherium paulhiacense*. However, if several taxa have been well studied at the locality rodents and lagomorphs: Werner 1994; suids: Hellmund 1991; carnivores: Heizmann and Morlo 1994; Peigné and Heizmann 2003; lizards: Klembara et al. 2017; all fauna: Heizmann et al. 1989; Costeur et al. 2012), it is not the case of large herbivores, and notably the rhinocerotids.

The locality of Ulm-Westtangente thus provides a unique opportunity to investigate rhinocerotids paleoecology at a key step of their evolutionary history. Here, we used a multi-proxy approach including diet proxies at different time scales (δ^13^C, first years of life; dental microwear, last days to months; and mesowear, general lifetime), habitat proxies (δ^18^O and δ^13^C: temperature, precipitation, habitat openness), life history proxies (enamel hypoplasia, body mass prediction, mortality curves: metabolic and environmental stresses, age structure, vulnerability periods). The combination of independent methods is intended to yield more robust results by compensating weaknesses of individual approaches, with the potential to shed light on new aspects and refine complex patterns. This will allow us to discuss niche partitioning, to propose paleoenvironmental interpretations and to provide new data for the understanding of the faunal turnover around the Oligocene-Miocene transition and the early diversification of the Rhinocerotidae in Europe.

### Abbreviations

Capital letters are used for the upper teeth (D: deciduous molar; P: premolar; M: molar), while lower case letters indicate lower teeth (d, p, m). Institutional abbreviations: SMNS – Staatliches Museum für Naturkunde Stuttgart, Germany.

## Materials and methods

The material studied is curated at the SMNS. It is composed of a total of 492 teeth: 337 of *Protaceratherium minutum* (137 isolated teeth, 200 from four skulls, two maxillae, 18 hemi-mandibles and mandibles, and 13 sets of associated isolated teeth) and 155 of *Mesaceratherium paulhiacense* (73 isolated teeth, 82 from two maxillae, seven hemi-mandibles and mandibles, and nine sets of associated isolated teeth; see Supplementary 1). The number of teeth studied with each method depended on the associated constraints (e.g., only averaged-worn upper molars with cusps preserved on ectoloph profile for mesowear) and is detailed thereafter (Table 1). Regarding isotopic content, the sample is very limited, as we focused on identifiable fragments (species and locus) because the method is destructive.

**Table 1:** Number of specimens studied by method and species of rhinocerotids from Ulm- Westtangente.

|  | *Protaceratherium minutum* | *Mesaceratherium paulhiacense* |
| --- | --- | --- |
| **Mesowear** | 16 | 12 |
| **Microwear** |  |  |
| **Grinding** | 21 | 11 |
| **Shearing** | 9 | 5 |
| **Enamel hypoplasia** | 317 | 149 |
| **Body mass** | 54 | 24 |
| **Stable isotopy ($\delta^{18}\text{O}$, $\delta^{13}\text{C}$)** | 3 | 5 |
| **Mortality curves** | 336 | 155 |

### Mesowear

Mesowear is the categorization of gross dental wear into herbivore diet categories. Traditionally, it is based on scoring cusp shape and occlusal relief on upper molars (Fortelius and Solounias 2000). Here, we used the microwear ruler developed by Mihlbachler et al. (2011) on horses (close relatives of rhinoceros), that gives scores from 0 (high sharp) to 6 (low blunt; see Supplementary 2 Fig 1). With this method browsers have low scores (attrition – tooth-tooth-contact - dominated) and grazers high ones (abrasion – tooth-diet-contact - dominated), while mixed-feeders have intermediate values. Contrary to most studies on mesowear, we consistently scored the paracone and not the sharpest buccal cusp (metacone or paracone), as important differences are noted between these two cusps in rhinoceros (Taylor et al. 2013; Hullot et al. 2021). Eventually, as mesowear score can be affected both by age and hypsodonty (Fortelius and Solounias 2000), we only scored upper molars with an average wear (wear stages from 4 to 7 defined by Hillman-Smith et al. 1986) and we compared the hypsodonty index (height of m3 divided by its width; Janis 1988) of both species.

**Figure 1:**
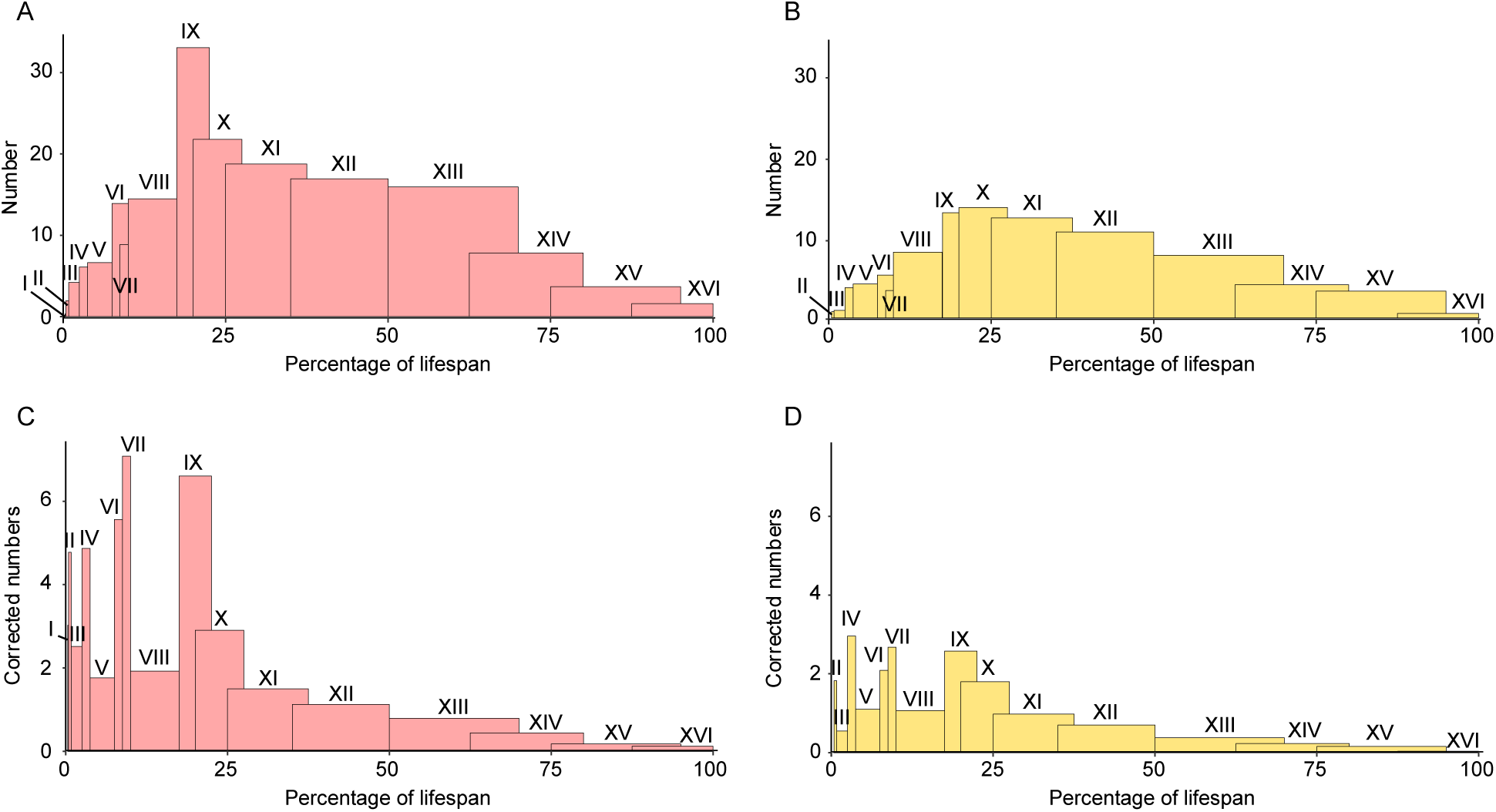
Mortality curves of both rhinocerotid species from Ulm-Westtangente. Number of specimens per age class by species: A – *Protaceratherium minutum*, and B – *Mesaceratherium paulhiacense* Correction for age class duration (number of specimens divided by the duration of the age class) by species: C – *Protaceratherium minutum*, and D – *Mesaceratherium paulhiacense* Classes expressed as percentage of lifespan instead of age to limit actualism Colour code: pink – *Protaceratherium minutum*, yellow – *Mesaceratherium paulhiacense*

### Dental Microwear Texture Analyses (DMTA)

Dental microwear texture analyses (DMTA) study dietary preferences at a short term scale (day to months; Hoffman et al. 2015). This technique examines tooth surface and identifies wear patterns associated with the different diet categories. In this study, we followed a protocol adapted from Scott et al. (2005) using scale-sensitive fractal analyses. We selected well-preserved molars (upper and lower, left and right) and sampled wear facets from both phases of the mastication (grinding and shearing) on the same enamel band near the protocone, protoconid or hypoconid (see Supplemetary 2 Fig 2). Facets were cleaned twice using a cotton swab soaked in acetone to remove dirt, grit and glue. Then two silicone molds (Coltene Whaledent PRESIDENT The Original Regular Body, ref. 60019939) were made. The second one was used for the analyses described hereafter.

**Figure 2:**
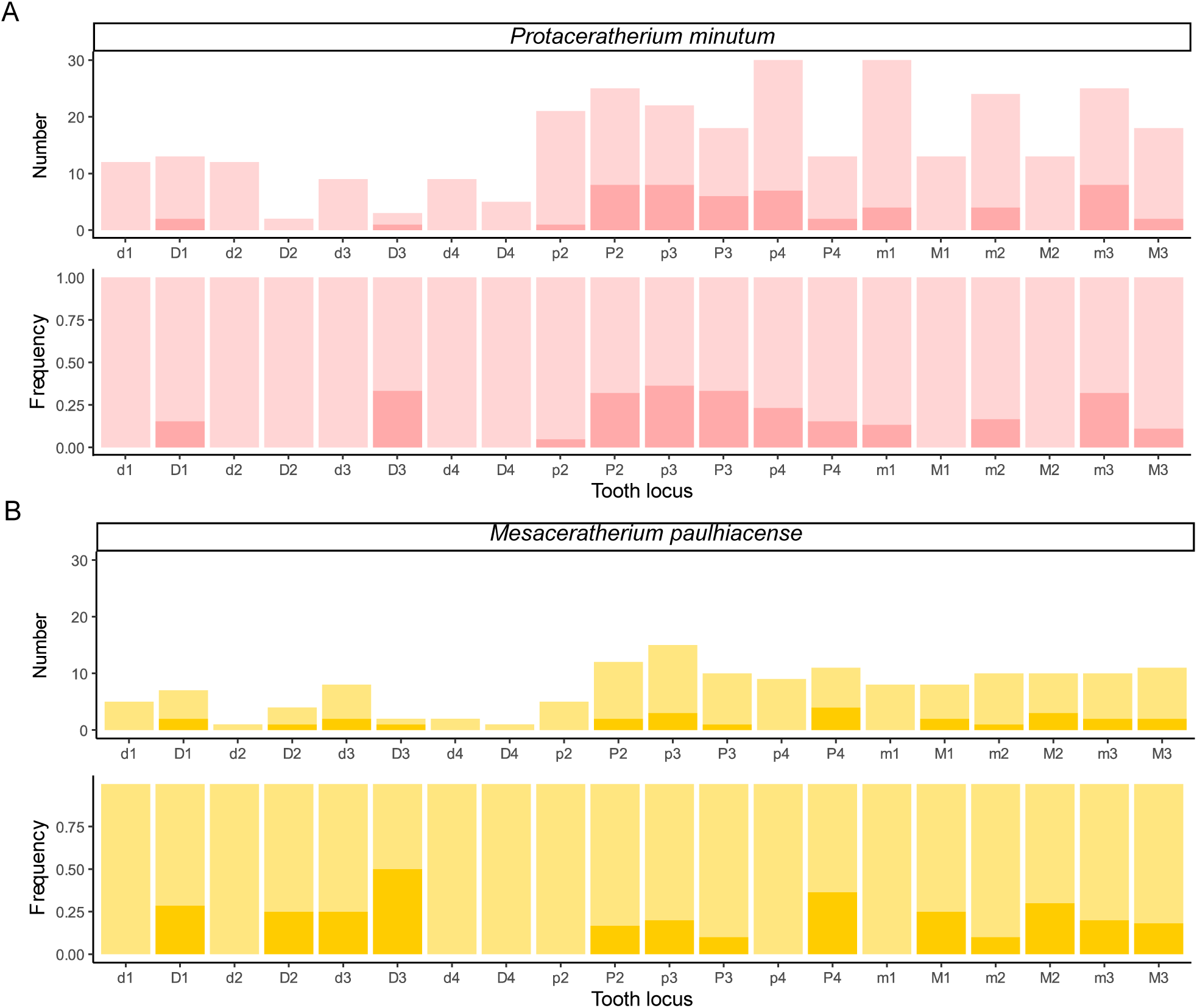
Number and frequency of hypoplasia by locus and species among the rhinocerotids from Ulm-Westtangente. Light colors for unaffected and dark for hypoplastic teeth A – Number and frequency of hypoplastic teeth vs. unaffected ones for *Protaceratherium minutum* B - Number and frequency of hypoplastic teeth vs. unaffected ones for *Mesaceratherium paulhiacense*

The facet was cut out of the mold, put flat under a Leica Map DCM8 profilometer (TRIDENT, PALEVOPRIM Poitiers) and scanned using white light confocal technology with a 100× objective (Leica Microsystems; Numerical aperture: 0.90; working distance: 0.9 mm). Using LeicaMap (v.8.2; Leica Microsystems), we pre-treated the obtained scans (.plu files) as follows: inversion of the surface (as they come from negative replica), replacement of the missing points (i.e, non-measured, less than 1%) by the mean of the neighboring points, removal of aberrant peaks (automatic operators including a morphological filter see supplementary Information in Merceron et al. (2016b), leveling of the surface, removal of form (polynomial of degree 8) and selection of a 200 × 200 μm area (1551 × 1551 pixels) saved as a digital elevation model (.sur) to be used for DMTA. We conducted scale-sensitive fractal analyses on the selected surfaces (200 × 200 μm; see Supplementary 3) using MountainsMaps® (v.8.2). Our study will focus on the following texture variables, described in detail in Scott et al. (2006):

- anisotropy or exact proportion of length-scale anisotropy of relief (epLsar) is a measure of the orientation concentration of surface roughness; in MountainsMaps®, this parameter has been corrected (NewepLsar), as there was an error in the code of Toothfrax (software previously used for DMTA but not supported anymore; Calandra et al. 2022);
- complexity or area-scale fractal complexity (Asfc) assesses the roughness at a given scale;
- heterogeneity of the complexity (HAsfc) gives information of the variation of complexity at a given scale (here 3 × 3 and 9 × 9) within the studied 200 × 200 μm zone.

### Body mass estimations

Body mass is linked to many physiological and ecological parameters (e.g., diet, metabolism, heat evacuation; Peters 1983; Owen-Smith 1988; Clauss et al. 2003). Many studies have established equations to estimate fossil body mass based on various dental and limb bone proxies (see the review of Hopkins 2018). Here, we chose to use several dental proxies, as we studied teeth for all other methods and as they are abundant and well preserved in the fossil record. Each dental proxy and its associated established equation is listed in Table 2 alongside with the corresponding references.

**Table 2:**
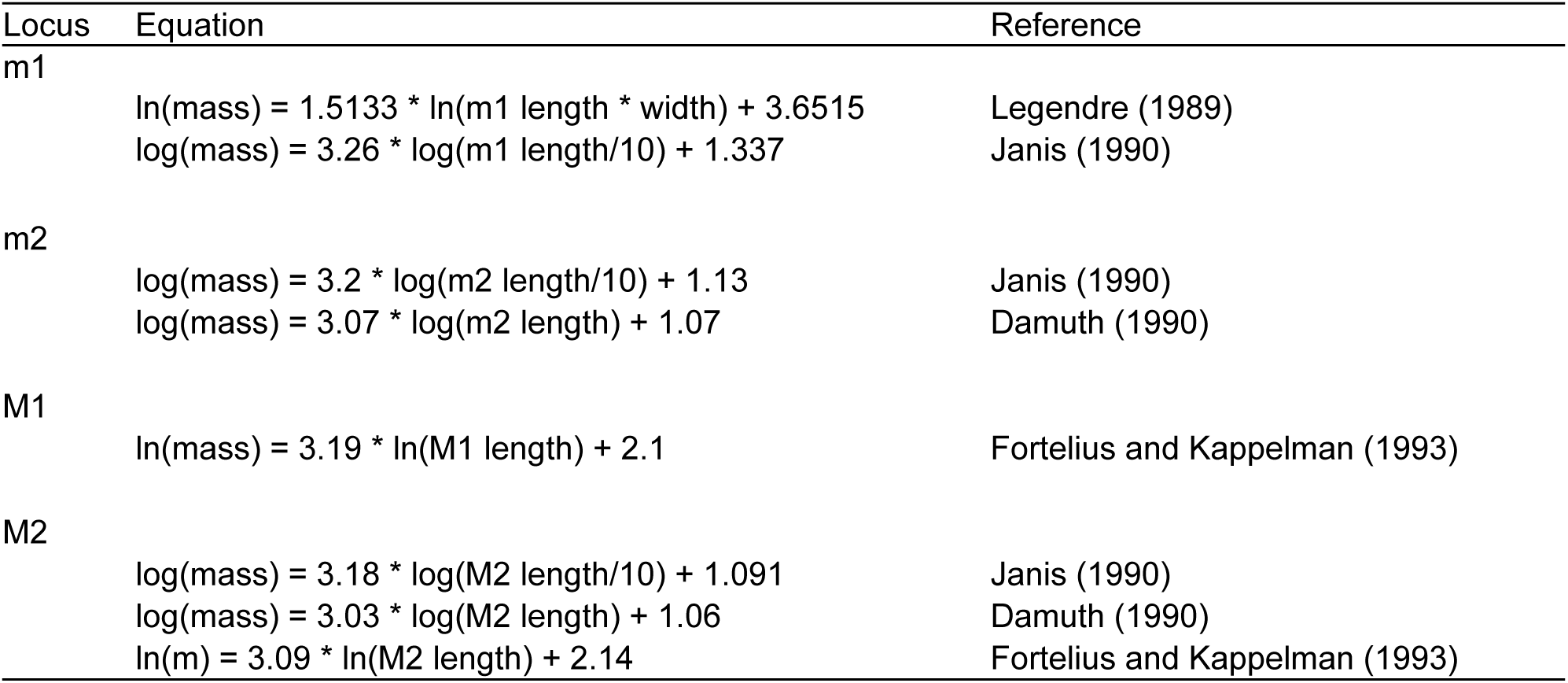
List of the dental proxies and the associated equations used to estimate body mass in this study. Measurements are in mm for all equations and give body mass in kg for Janis (1990) and in g otherwise.

### Hypoplasia

Hypoplasia is a permanent and sensitive defect of the enamel that has been correlated with stresses, notably environmental (Skinner and Pruetz 2012; Kierdorf et al. 2012; Upex and Dobney 2012). It is however non specific and can take several forms, the etiology of which is not well understood (Small and Murray 1978). In the literature, no standard protocol, nor threshold between normal and pathological enamel are available, so we followed a classical approach that consists in recording and categorizing the defects following the *Fédération Dentaire Internationale* (1982). This index recognizes three main types of defects: linear (line around the crown), pitted (restricted rounded defect) and aplasia (extended zone missing enamel; see Supplemetary 2 Fig 3). In parallel, we recorded several parameters (distance to enamel-dentin junction, width if applicable, localization on the tooth, severity). We studied all available identified cheek teeth, both deciduous and permanent, with the exception of very worn (wear stage 9 and 10 defined by Hillman-Smith et al. 1986) or damaged teeth (i.e., with limited to no enamel preserved) to limit the risk of false negatives. This resulted in the exclusion of 26 teeth (20 of *P. minutum* and six of *M. paulhiacense*) and left 466 teeth to study.

**Figure 3:**
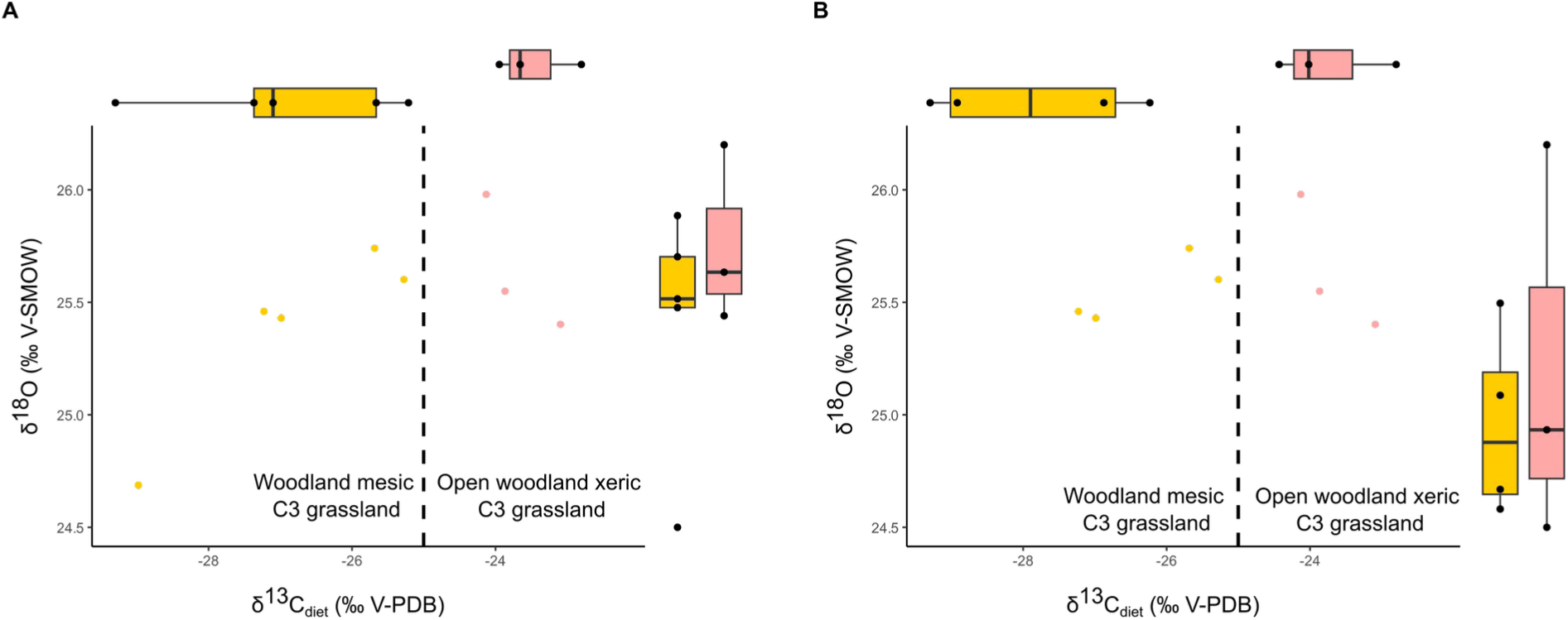
Isotopic values of δ^18^OCO3, SMOW and δ^13^Cdiet for the two rhinocerotids from Ulm-Westtangente. A-Dotplot of δ^18^OCO3, SMOW against δ^13^Cdiet (corrected for body mass and for the variations of atmospheric δ^13^CCo2) with associated boxplots B-Same graph without the outlier of *Mesaceratherium paulhiacense* One specimen of *M, paulhiacense* appeared as an outlier for δ^18^OCO3, SMOW (outside whiskers range) and was removed as we suspected a weaning signal Color code: pink-*Protaceratherium minutum* and yellow-*Mesaceratherium paulhiacense* Threshold for modern plants and environment reported in Domingo et al. (2013).

### Mortality curves and age structure

Mortality curves are graphs indicating the number of individuals from a sample dying at each age category. They are often used to study taphonomy (attritional causes) and to infer population structure in ancient communities (Fernandez and Legendre 2003; Bacon et al. 2018). Here, we used a protocol specifically designed for rhinocerotids detailed in Hullot and Antoine (2020). Age estimation is based on the correspondence between wear stages (1-10) and age classes (0-XVI) defined by Hillman-Smith et al. (1986) for each tooth locus. Mortality curves can then be established by following the steps hereunder:

- each tooth is considered individually and has a weight of 1;
- estimation of the wear stage (1 to 10) for each tooth;
- correlation to one or several age classes corresponding to the wear observed for the locus concerned;
- equal weight given to each age class (1 if one, ½ if two, and so on);
- for associated teeth, grouping of all teeth as a single individual and proposition of a class or combination of weighted classes for the group;
- construction of mortality curves from the weighted classes.

Based on the individual ages estimated following the steps above, we calculated the age structure of our sample. Ontogenetic stages are defined as follows, based on the correlations from Hullot and Antoine (2020). Juveniles (birth to weaning) include age classes from I to V, corresponding to 1.5 months to 3 years in the extant white rhinoceros and ending with the eruption of the first permanent teeth (m1/M1). Subadults (weaning to sexual maturity) correspond to age classes VI to VIII (i.e., 3 to 7 years), ending with the eruption of the last permanent teeth (m3/M3). Adults (sexual maturity to death) correlates with age classes IX to XVI (i.e., 7 to 40 years), after the eruption of the last permanent teeth (m3/M3).

### Minimum number of individuals

The minimum number of individuals (MNI) is the smallest number of individuals of the same species that can be identified from a fossil assemblage. It is determined by the count of the most abundant anatomical element from the same side (e.g., left femurs, right M3s). Here, estimations also took into account the incompatibility groups based on pattern of dental eruption defined by Hullot and Antoine (2020): for instance milk teeth are never associated in a functioning tooth row with fourth premolars (group b) nor third molars (group a).

### Isotopic analyses

Stables isotopes are often used in paleontology as they allow for dietary and environmental insights into terrestrial and aquatic ecosystems (Cerling et al. 1997; Clementz 2012). Here, we studied δ^13^C – linked to the feeding behavior (C3-C4 plants) and to the habitat openness – and δ^18^O – that depends on temperature and precipitation – of the carbonates from dental enamel of both rhinocerotid species from Ulm- Westtangente. The δ^18^O of rhinoceros enamel is especially interesting for climatic/environmental reconstructions as the animals were abundant and large sized, with a drinking behavior likely resulting in a δ^18^O accurately recording the meteoritic precipitation (Clauss et al. 2005; Levin et al. 2006; Martin et al. 2008; Zanazzi et al. 2022).

After mechanical cleaning of a small zone, samples were taken on identified tooth fragments (taxon and locus), preferably from third molars to avoid pre-weaning or weaning signal, using a Dremmel© equipped with a diamond tip. Between 500 and 1000 μg were then used for the analyses. Organic matter was removed following standard procedures (Cerling et al. 1997) and the samples were then acidified with phosphoric acid (103 %), producing carbon monoxide to be analyzed for isotopic content using a Micromass Optima Isotope Ratio Mass Spectrometer (Géosciences Montpellier). Results (Table 3) are expressed as ratio (‰) to the Vienna-Pee Dee Belemnite standard as follows:

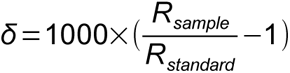

**Table 3:** O and C isotopic compositions of the enamel carbonates of Ulm-Westtangente rhinocerotids.

| Specimen | Weight (mg) | Tooth | Species | $\delta^{13}\text{C}_{\text{CO}_3, \text{enamel}}$ (‰ V-PDB) | $\delta^{13}\text{C}_{\text{CO}_3, \text{enamel}}$ Stdev | $\delta^{18}\text{O}_{\text{CO}_3, \text{enamel}}$ (‰ V-PDB) | $\delta^{18}\text{O}_{\text{CO}_3, \text{enamel}}$ Stdev |
| --- | --- | --- | --- | --- | --- | --- | --- |
| 47950 | 628 | M3 | *P. minutum* | -7.47 | 0.01 | -5.34 | 0.04 |
| 48079 | 520 | M3 | *P. minutum* | -8.25 | 0.02 | -5.2 | 0.03 |
| 48184 | 580 | M3 | *P. minutum* | -8.51 | 0.04 | -4.78 | 0.06 |
| 48072 | 628 | m3 | *M. paulhiacense* | -9.06 | 0.02 | -5.15 | 0.04 |
| 48008 | 722 | M1 | *M. paulhiacense* | -10.77 | 0.02 | -5.32 | 0.03 |
| 48183 | 502 | P4 | *M. paulhiacense* | -12.76 | 0.01 | -6.04 | 0.02 |
| 47877 | 1054 | M3 | *M. paulhiacense* | -11.01 | 0.02 | -5.29 | 0.02 |
| 47877 | 554 | M1-2 | *M. paulhiacense* | -9.47 | 0.01 | -5.01 | 0.02 |

where R_sample_ refers to the ratio of 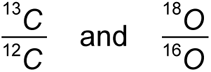 of the sample and R_standard_ to the Vienna-Pee Dee Belemnite standard. The within-run precision (± 1 σ) of these analyses as determined by the replicate analyses of NBS 18 and AIEA-603 was less than ± 0.2 ‰ for δ^13^C and ± 0.3 ‰ for δ^18^O (n = 5-6 respectively).

The δ^13^C_diet_ can be traced back from δ^13^C_CO3, enamel_ taking into account the body mass and the digestive system (Tejada-Lara et al. 2018) as detailed below:

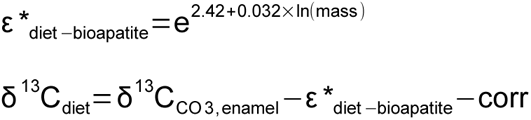

where corr is the correction factor for the variation of δ^13^C_CO2_ of the atmosphere, here equal to 1.9 ‰. Post 1930, the value of δ^13^C_CO2_ in the atmosphere is -8 ‰ (Zachos et al. 2001). For Ulm-Westtangente, the reconstructed values based on benthic foraminifera at around 22 Mya (Tipple et al. 2010) are higher than today, with an estimate of -6.1 ‰.

The δ^13^C_diet_ can in turn provide the mean annual precipitation (MAP) with the equation from Kohn (2010):

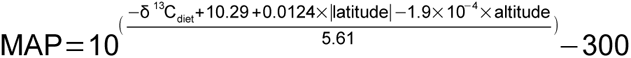

The δ^18^O_Co3(V-PDB)_ was converted into δ^18^O_Co3(V-SMOW)_ using the equation from Coplen et al. (1983):

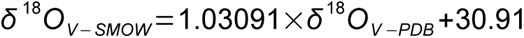

This was used to calculate the δ^18^O_precipitation_ and the mean annual temperature (MAT) detailed as follows. No reliable equation to estimate the δ^18^O_precipitation_ based on the δ^18^O_enamel_ of rhinoceros is available in the literature. Tütken et al. (2006) tentatively established one, based on a dataset of extant rhinoceros (including zoo specimens) and using converted phosphates values obtained from the carbonates ones. The MATs obtained with this equation are consistently 2-4°C higher (MH pers. obs.) than the ones estimated with equations specific to horses (Sánchez Chillón et al. 1994), bisons (Bernard et al. 2009), or elephants (Ayliffe et al. 1992). Hence, we decided to use the equation for modern elephants, as they might be the closest equivalent to rhinoceros for isotope fractionation. The δ^18^O in the following equations are expressed in relation to the V-SMOW:

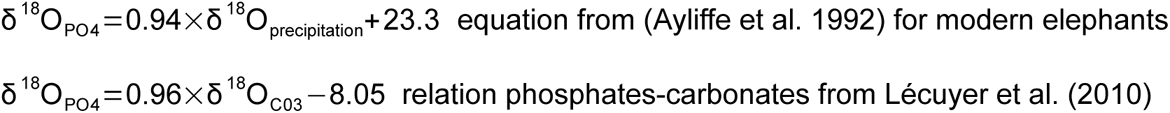

Hence: δ^18^O_precipitation_ 18 = 1.02×δ^18^O_co3_ −33.3

Eventually, the MAT (Tütken et al. 2006) can be calculated using the obtained δ^18^O_precipitation_:

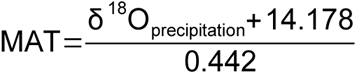

### Statistics and Figures

All statistics were conducted in R (R Core Team, 2021: v. 4.1.2) equipped with the package tidyr (Wickham and Henry 2020), MASS (Ripley et al. 2013) and mvnormtest (Jarek, 2012). Following the recent statement of the American Statistical Association (ASA) on p-values (Wasserstein and Lazar 2016), we favored giving exact values and we tried to be critical regarding the classical thresholds of “statistical significativity”. Figures were done using R packages ggplot2 (Wickham 2016), cowplot (Wilke 2020), as well as Inkscape (v. 1.0.1).

## Results

### Structure of the rhinocerotid sample from Ulm-Westtangente

There are about twice as many teeth attributed to *Protaceratherium minutum* (337) than to *Mesaceratherium paulhiacense* (155) at Ulm-Westtangente. The minimum number of individuals (MNI), based on dental remains, is 17 for *P. minutum* (number of left m1) and 10 for *M. paulhiacense* (number of left p3). When dental eruption incompatibility groups are considered, the MNIs are 24 (group b: fourth premolars and milk teeth) and 15 (group d: third milk molars and third premolars) respectively.

Before correction for the duration of the age classes, we observed a single peak centered around age class IX (Figure 1A-B) for both species, although more spread out in *M. paulhiacense* (IX to XII). In the corrected mortality curves (i.e., number at each age class divided by the duration of the age class), similar tendencies were observed for both species (two-sided Kolmogorov-Smirnov test: D = 0.375, p-value = 0.2145), but with a smaller amplitude for the less abundant *M. paulhiacense*. The histograms of both species revealed four distinct peaks, around age classes I-II, IV, VI-VII, and IX (Figure 1C-D).

The age structure of our rhinocerotid sample from Ulm-Westtangente both species merged is composed by 10.8 % of juveniles, 20.2 % of subadults and 69 % of adults. The proportions were similar between both species (Chi2: X-squared = 0.32999, df = 2, p-value = 0.8479), with 10.5 % of juveniles, 18.3 % of subadults and 71.2 % of adults for *M. paulhiacense*, and 11 % of juveniles, 21.2 % of subadults and 67.8 % of adults for *P. minutum* (see details in Supplementary 1).

### Enamel hypoplasia

At Ulm-Westtangente, a total of 79 teeth out of the 466 studied for hypoplasia (16.95 %) bear at least one hypoplastic defect. There are however frequency discrepancies depending on species and tooth loci (Figure 2).

Both rhinocerotid species have similar overall hypoplasia prevalences (Kruskal-Wallis test: chi-squared = 0.044163, df = 1, p-value = 0.8336) with 16.7 % (53/317) for *P. minutum* and 17.4 % (26/149) for *M. paulhiacense*. Regarding tooth loci, when both species are merged, four are not affected (only milk molars), 14 have a prevalence of hypoplasia above 10 % and seven above 20 % (Table 4). In general, milk teeth are relatively spared (11.7 to 20 % for D1, D2 and d3) or even not affected (0 % of d1, d2, d4, and D4), with the notable exception of D3 for both species (species merged: 3/5, 40 %), although the sample is very limited (Table 4).

**Table 4:** Hypoplasia prevalence by locus (both species merged) at Ulm-Westtangente.

| Locus | d1 | D1 | d2 | D2 | d3 | D3 | d4 | D4 | p2 | P2 | p3 | P3 | p4 | P4 | m1 | M1 | m2 | M2 | m3 | M3 |
| --- | --- | --- | --- | --- | --- | --- | --- | --- | --- | --- | --- | --- | --- | --- | --- | --- | --- | --- | --- | --- |
| Unaffected | 17 | 16 | 13 | 5 | 15 | 3 | 11 | 6 | 25 | 27 | 26 | 21 | 32 | 18 | 34 | 19 | 29 | 20 | 25 | 25 |
| Affected | 0 | 4 | 0 | 1 | 2 | 2 | 0 | 0 | 1 | 10 | 11 | 7 | 7 | 6 | 4 | 2 | 5 | 3 | 10 | 4 |
| Frequency | 0 | 20 | 0 | 16.67 | 11.76 | 40 | 0 | 0 | 3.85 | 27.03 | 29.73 | 25 | 17.95 | 25 | 10.53 | 9.52 | 14.71 | 13.04 | 28.57 | 13.79 |

The pattern of loci affected by hypoplasia was however different by species. For *P. minutum*, the most affected teeth were p3 (8/22; 36.4 %), D3 (1/3; 33.3 %), P3 (6/18; 33.3 %), P2 and m3 (each 8/25; 32 %), while d1, d2, D2, d3, d4, D4, M1, and M2 were never hypoplastic (Figure 2). On the other hand, in *M. paulhiacense* the most affected teeth were D3 (1/2; 50 %), P4 (4/11; 36.4 %), M2 (3/10; 30 %), D1 (2/7; 28.6 %), D2 (1/4; 25 %), d3 (2/8; 25 %) and M1 (2/8; 25 %), while d1, d2, d4, D4, p2, p4, and m1 were never hypoplastic (Figure 2).

### Body mass

Both species exhibit obvious differences in size, which is reflected in their body mass estimates, as detailed in Table 5. Mean estimates range from 438 to 685 kg for *P. minutum* and from 1389 to 2327 kg for *M. paulhiacense*, depending on the dental proxy and the equation used.

**Table 5:**
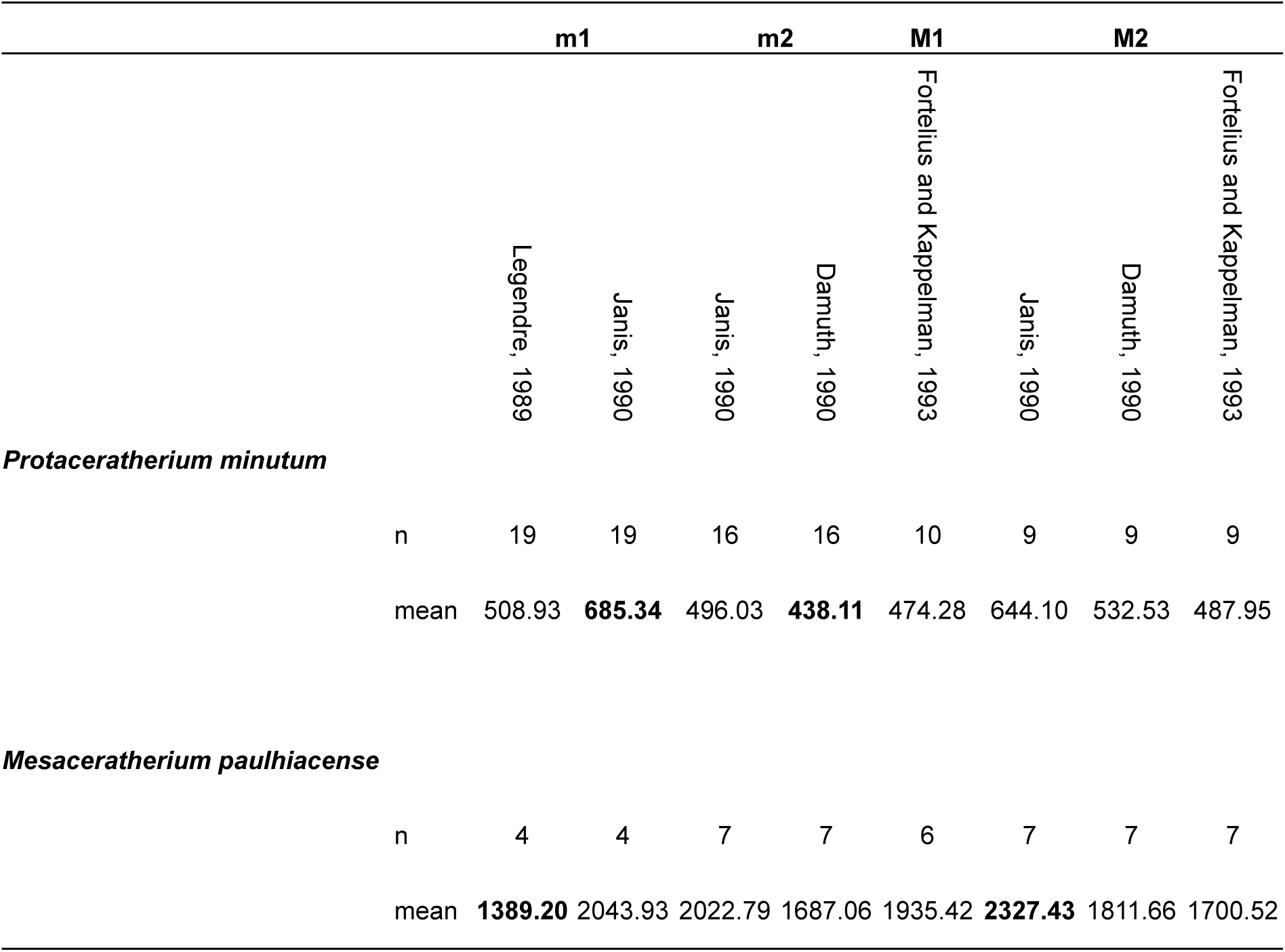
Body mass estimates (in kg) of the rhinocerotids from Ulm-Westtangente based on dental proxies. The number of teeth (n; only one per specimen if both were available) and the mean are shown for each proxy and species. Minimum and maximum estimates are indicated in bold for each species.

### Dietary preferences and habitat

Only a few unworn m3 were available to calculate the hypsodonty index (n = 5 for *P. minutum* and n = 3 for *M. paulhiacense*), but both species had relatively similar values (Kruskal-Wallis chi-squared = 1.0889, df = 1, p-value = 0.2967). However, according to the thresholds established by Janis (1988), *P. minutum* classifies as brachyodont (mean hypsodonty index of 1.31), while *M. paulhiacense* is mesodont (mean hypsodonty index of 1.55).

The mesowear scores were contrasted between the two species samples (Table 6; Kruskal-Wallis test: chi-squared = 5.2828, df = 1, p-value = 0.02154). *Protaceratherium minutum* (n = 16) had a mean ruler score of 1.75, with values ranging from 0 (only one tooth) to 3 (only one tooth). On the other hand, *Mesaceratherium paulhiacense* (n = 12) had a higher mean ruler score of 2.25 with values ranging from 1 (only two teeth) to 4 (five teeth). This suggests a more abrasive diet for *M. paulhiacense* consistent with the slightly higher hypsodonty index obtained above for this species (mesodont).

**Table 6:** Dental wear signatures of the rhinocerotids from Ulm-Westtangente. When several teeth per specimen were available, only one tooth per specimen was studied, preferentially second molars. Gr: grinding facet, Sh: shearing facet, sd: standard deviation

|  | *Protaceratherium minutum* |  |  | *Mesaceratherium paulhiacense* |  |  |
| --- | --- | --- | --- | --- | --- | --- |
|  | n | mean | sd | n | mean | sd |
| **Mesowear** | 16 | 1.75 | 0.68 | 12 | 2.25 | 0.97 |
| **Microwear** | Gr: 21<br>Sh: 9 |  |  | Gr: 11<br>Sh: 5 |  |  |
| **NewepLsar ($10^{-2}$)** |  | Gr: 1.67<br>Sh: 1.84 | Gr: 0.17<br>Sh: 0.28 |  | Gr: 1.70<br>Sh: 1.88 | Gr: 0.12<br>Sh: 0.19 |
| **Asfc** |  | Gr: 1.72<br>Sh: 1.02 | Gr: 1.25<br>Sh: 0.67 |  | Gr: 1.32<br>Sh: 0.77 | Gr: 0.71<br>Sh: 0.22 |
| **HAsfc9** |  | Gr: 0.37<br>Sh: 0.29 | Gr: 0.44<br>Sh: 0.14 |  | Gr: 0.24<br>Sh: 0.35 | Gr: 0.07<br>Sh: 0.16 |
| **HAsfc81** |  | Gr: 0.68<br>Sh: 0.46 | Gr: 0.60<br>Sh: 0.13 |  | Gr: 0.40<br>Sh: 0.58 | Gr: 0.10<br>Sh: 0.25 |

The dental microwear texture analyses (DMTA) revealed slightly contrasted results between grinding and shearing facets (MANOVA: p-value = 0.07842), but only for NewepLsar (ANOVA, p-value = 0.00621) and Asfc (ANOVA, p-value = 0.041). Regarding species, DMT signatures were relatively similar (Table 6; MANOVA: p-value = 0.65653), although some differences might be worth noting. The mean Asfc was lower for both facets of *M. paulhiacense* (Gr: 1.32; Sh: 0.77) than that of *P. minutum* (Gr: 1.72; Sh: 1.02), suggesting the processing of slightly softer food items *for M. paulhiacense*. On the grinding facet, *P. minutum* had higher mean values of HAsfc9 and HAsfc81 (0.37 and 0.68) than *M. paulhiacense* (0.24 and 0.40), but the opposite is observed for the shearing facet (HAsfc9: 0.29 for *P. minutum* vs 0.35 for *M. paulhiacense*; HAsfc81: 0.46 vs 0.58 respectively). This indicates a greater diversity of items consumed requiring grinding and a lower diversity of items consumed requiring shearing in *P. minutum*.

The isotopic content of the enamel carbonates for our rhinocerotid sample (n = 8) ranged between -12.76 and -7.47 ‰ for δ^13^C_CO3, enamel_ and -6.04 and -4.78 for δ^18^O_CO3, enamel_ (Table 3). Regarding δ^18^O_CO3, enamel_, both rhinocerotids had similar means (Wilcoxon test; W = 6, p-value = 0.7857) with -5.1 ± 0.3 ‰ Vienna Pee Dee Belemnite (VPDB) for *P. minutum* and -5.4 ± 0.4 ‰ VPDB for *M. paulhiacense*, suggesting similar drinking water sources. The mean δ^18^O_CO3, enamel_ was however a little lower for *M. paulhiacense*, but this could be due to an outlier for this species (Figure 3A; mean without outlier -5.2 ± 0.1 ‰ VPDB). The δ^18^O_precipitation_ obtained from δ^18^O_SMOW_ had a mean of -7.2 ± 0.2 ‰ (both species merged, outlier removed), which is significantly higher than today’s values that are around -9.3 ‰ (today’s value at Ulm, Germany calculated with the Online Isotopes in Precipitation Calculator: https://wateriso.utah.edu/waterisotopes/pages/data_access/oipc.html). The value obtained from the rhinocerotid specimens (both species merged, outlier removed) suggests a mean annual temperature (MAT) of 15.8°C (see Supplementary 1 for all calculated values for each specimen).

Clear differences were observed between both species for δ^13^C_diet_ (Wilcoxon test; W = 0, p-value = 0.03571) although they stay in the range of C3 feeding and habitats (Figure 3): *P. minutum* had higher values (n = 3, mean = -23.7 ± 0.5 ‰) compared to that of *M. paulhiacense* (n = 5, mean = -26.8 ± 1.5 ‰). Without the outlier (*M. paulhiacense*, mean = -26.3 ± 1 ‰), the difference between the two species for δ^13^C_diet_ is less marked (Wilcoxon test; W = 0, p-value = 0.05714), but still suggests different ecologies (diet and/or habitat). From the δ^13^C_diet_ we calculated the mean annual precipitation (MAP). All specimens of *P. minutum* yielded negative MAPs, while the mean for *M. paulhiacense* (outlier excluded) gave a MAP of 317 mm/year (see Supplementary 1). The MAP calculated based on *M. paulhiacense* however showed great variations depending of the specimen (84 to 556 mm/year).

## Discussion

### Body mass of the rhinocerotids from Ulm-Westtangente

In extant rhinoceros, one of the most accurate dental proxy appears to be the length of M2 with the equation of Damuth (1990; MH pers. obs.), which here gives means of 532.5 kg for *P. minutum* and 1811.7 kg for *M. paulhiacense* (Table 5). Such a body mass for *P. minutum*, around 500 kg, is consistent with a previous estimation based on the astragalus (Becker 2003). In their study on Ulm-Westtangente, Costeur et al. (2012) used the occlusal surface of m1 as a proxy for body mass (Legendre 1989). They found higher means than our study using the same proxy for both species (590 vs. 509 kg and 1752 vs. 1389 kg respectively). Such discrepancies might result from an inter-observer bias, the inclusion of misidentified teeth (e.g., m2 instead of m1), a restricted data set in their study (sampling bias), or the inclusion of worn teeth in our study (rhinoceros m1 tend to shorten with wear; Antoine pers. comm., MH pers. obs.). In any case, the rhinocerotids are the biggest species found at Ulm-Westtangente (Costeur et al. 2012), with only *M. paulhiacense* exceeding 1000 kg, the threshold for megaherbivores following Owen-Smith (1988). Such a result is not really surprising, as European faunas did not show a high diversity of megaherbivores before the late early Miocene (Proboscidean Datum Event; Costeur et al. 2012).

### Age structure, stress vulnerability and mortality of the rhinocerotids from Ulm-Westtangente

The age structure with around 10 % of juveniles, 20 % of subadults and 70 % of adults for both rhinocerotid samples is close to the structure observed in modern *Diceros bicornis* populations (Supplementary 2 Fig 4; Goddard 1970; Hitchins 1978). As Ulm-Westtangente provides a relatively time constrained framework (fossil layer only 35 cm thick; Heizmann et al. 1989), the studied sample could closely reflect living populations of both rhinocerotids with very minor preservation or taphonomic bias. The comparison with extant populations of rhinoceros must, however, be drawn with caution, as poaching and population management might have biased the observed age structure.

The mortality curves based on rhinocerotid teeth from Ulm-Westtangente (Figure 1) suggested several vulnerability periods, similar in both species. The first peak is noted for classes I and II (0.31 to 0.83 % of lifespan, i.e. 1.5 to 4 months old in extant *Ceratotherium simum*; Hillman-Smith et al. 1986), and corresponds to the period shortly after birth. Birth is known to be a particularly stressful event in the life of an animal (Upex and Dobney 2012), and has notably been correlated with the presence of a neonatal line in primate deciduous teeth and first molar (Gustafson and Gustafson 1967; Risnes 1998), as well as with hypoplasia on fourth milk molars in several ungulates (rhinoceros: Mead 1999; bison: Niven et al. 2004; sheep: Upex and Dobney 2012). In our data set d4/D4 were never affected in either species (Figure 2). This finding is not necessarily contradicting, as the animal must survive the stress for the hypoplasia defect to be visible (Guatelli-Steinberg 2001). Other loci, that also develop around birth, displayed however a higher prevalence of hypoplasia, especially in *M. paulhiacense*. This is notably the case of the first molar (Tafforeau et al. 2007; Upex and Dobney 2012), which was amongst the most affected loci in *M. paulhiacense* (25 % of M1; Figure 2), or of other milk teeth loci suggesting early life stresses (or even *in utero*; Figure 2).

The second peak is observed at age class IV (2.5 to 3.75 % of lifespan, i.e. 1 to 1.5 years old), which could be a sign of juvenile diseases. Hypoplasia in fossil rhinoceros has previously been linked to juvenile malnutrition or disease causing a fever (Bratlund 1999), although no specific locus was mentioned. Based on the timing of tooth development in extant rhinoceros, hypoplasia due to juvenile diseases could impact all premolars and molars but m3/M3 (chapter IV, page 131: Hullot, 2021). All these loci are affected in our data set, with a variable prevalence depending on the species, but a direct correlation with juvenile diseases is not straight forward.

The third peak corresponds to age classes VI-VII (7.5 to 10 % of lifespan, i.e. 3 to 4 years old), which are correlated to the period shortly after weaning (Hullot and Antoine 2020), maybe indicating cow-calf separation. Weaning and cow-calf separation are known to be critical times for many extant mammals, including rhinoceroses. Indeed, at that time, adult size is not yet reached (Owen-Smith 1988), leading to a higher predation risk (Brain et al. 1999). In parallel, calves might experience food stresses (Mead 1999) due to their new independence. Interestingly, both events have been associated with hypoplastic defects in primates and pigs (Goodman and Rose 1990; Dobney and Ervynck 2000; Guatelli-Steinberg 2001; Skinner and Pruetz 2012). In fossil rhinoceroses, Mead (1999) supposed that hypoplasia on the fourth premolars might be associated with cow-calf separation. Here, fourth premolars are mildly to highly affected (15 to 36 %) for both species (Figure 2, Table 4), which could suggest that cow-calf separation was also a stressful event in these fossil species. Moreover, in modern rhinoceroses, second molars develop at a relatively similar timing than that of fourth premolars (Goddard 1970; Hitchins 1978; Hillman-Smith et al. 1986). Interestingly, P4 and M2 commonly bear at least one hypoplastic defect in *M. paulhiacense* (Figure 2).

Eventually, a last peak in mortality is observed around classes IX-X (17.5 to 27.5 % of lifespan, i.e. 7 to 11 years old), correlating with sexual maturity (Hullot and Antoine 2020). In modern rhinoceroses, courtship and mating can be violent, including male fights for dominance, male rejection by the female, female chasing by males or mating wounds (Owen-Smith 1988; Dinerstein 2003), all of which could explain the increase of mortality observed for these age classes. However, potential stresses associated with this last peak were not recorded by enamel hypoplasia, as they occurred post-odontogenetically.

### Paleoecology of the rhinocerotids from Ulm-Westtangente

Overall, results from carbon isotopic content, meso- and micro-wear suggest different dietary preferences for the two rhinocerotid species at Ulm-Westtangente. The carbon isotopic signal of both species falls in the range of C3 feeding (modern C3 ranging from -20 to -37 ‰; Kohn, 2010) but points towards different habitats and/or dietary preferences (Figure 3). Based on the thresholds between habitats established by Domingo et al. (2013), the δ^13^C_diet_ values corrected for body mass and atmospheric δ^13^C_CO2_ variations (see material and methods) indicate a more closed environment (woodland mesic C3 grassland) for *M. paulhiacense*, whereas *P. minutum* seems restricted to open woodland xeric C3 grassland (Figure 3). This last finding is in line with what has been inferred based on morphology, as *Protaceratherium minutum* is described as a cursorial brachyodont species typical of forested, partially open environments (Becker 2003).

Regarding mesowear, *M. paulhiacense* has a higher mean score (2.25) than *P. minutum* (1.75), pointing towards a more abrasive diet (Table 6). Interestingly, *M. paulhiacense* has a slightly higher hypsodonty index and is also bigger (Table 5), which is often associated to a supposedly higher tolerance to low quality, more fibrous and abrasive diet (Jarman-Bell principle; Clauss et al. 2013; Steuer et al. 2014). The mesowear values observed for *P. minutum* (mostly 1 or 2) are consistent with browsing but also overlap with mixed-feeding, whereas that for *M. paulhiacense* (2, 3 and 4) point towards mixed-feeding or grazing (Rivals et al. 2017). However, a C4-grazing is unlikely based on the isotopic signal (Figure 3), the age and the situation of Ulm-Westtangente (C4 scarce or absent in the early Miocene of Germany; Strömberg, 2011). Fewer differences were noted in the microwear pattern of both species (Table 6). The slight differences in DMT are rather consistent with the mesowear, and points towards browsing in *P. minutum* (high values of Asfc correlated with browsing; Scott et al. 2006), and mixed-feeding in *M. paulhiacense*.

### Outlier in the stable isotope dataset: weaning signal?

One tooth of *M. paulhiacense* has significantly lower values for both δ^13^C_C,enamel_ and δ^18^O_C,enamel_ (Table 3) than the rest of the sampled teeth and, hence, was considered as an outlier. Interestingly, this is the only premolar of our data set (P4). Such different isotopic contents could be the result of a sampling in the pre-weaning part of the P4, as this tooth is known to partly develop before weaning in several extant rhinoceros species (Goddard 1970; Hitchins 1978; Hillman-Smith et al. 1986; Mead 1999). Indeed, milk has a higher δ^18^O than drinking water due to the preferential loss of the light oxygen isotope (^16^O) through expired and transcutaneous water vapor fluxes (Kohn et al. 1996), as well as a lower δ^13^C than plants due to the presence of lipids, which are depleted in ^13^C relatively to other macronutrients (DeNiro and Epstein 1977). The carbon depletion might however be relatively limited in the case of rhinoceros, as the milk of living individuals has a very low lipid content (Osthoff et al. 2021).

### Associated fauna and paleoenvironment

Besides the two rhinocerotids species, the herbivore assemblage at Ulm-Westtangente includes two other perissodactyl species – a chalicothere (cf. *Metaschizotherium wetzleri*) and a tapir (*Paratapirus intermedius*) – five artiodactyl species and 18 rodent and lagomorph species (Heizmann et al. 1989; Costeur et al. 2012). However, the body mass of all these species is significantly lower than that of the rhinocerotids, with only the chalicothere and the tapir ranging between 100 and 200 kg (Costeur et al. 2012), which limits potential competitive interactions. Unfortunately, no precise data is available on the paleoecology of these species, but schizotheriines are often reconstructed as open woodland dwellers (Heissig 1999) consuming leaves, fruits and maybe seeds and bark (Schulz et al. 2007; Semprebon et al. 2011) – which is relatively similar to our results for rhinocerotids – while tapirs prefer forested habitats and are typically folivores (DeSantis and MacFadden 2007).

Regarding environmental conditions, the MAP was around 317 mm/year suggesting rather dry conditions (Supplementary 1). Important individual variations were observed, which might point towards a certain seasonality. However, our sample is limited and restricted to a single taxon (Rhinocerotidae), and robust estimates of MAP require averaging over multiple species in a single locality (Kohn 2010). Moreover, *P. minutum* specimens yielded negative values of MAP. Some parameters might result in low to negative MAP, such as the consumption of C4 or the variation of C3 plant isotope compositions within a single locality (Kohn 2010).

Regarding temperature, the study of Costeur et al. (2012), has taken interest in the whole mammal fauna and proposed a cenogram. This study inferred an environment of a warm-temperate forest with grassland habitats at Ulm-Westtangente, but found quite a low MAT, around 7°C. This MAT is similar to the one proposed for Montaigu-le-Blin (France; reference of MN2a) using the same approach. The authors note that this estimation is surprising, as the Aquitanian conditions are reconstructed as warm-temperate to subtropical (Zachos et al. 2001), and argue that this could be a remnant of the Mi-1 glaciation. Here, we found a MAT of about 16°C based on the oxygen content of the enamel from our rhinocerotid sample. Although several complications exist in calculating absolute MAT from δ^18^O_precipitation_ (see details in Zanazzi et al. 2022), this result is more consistent with the warm-temperate forest with grassland habitats inferred on species assemblage and cenograms (Costeur et al. 2012), the presence of ectothermic species (Costeur et al. 2012; Klembara et al. 2017), and with the global climate reconstructed at that time (Zachos et al. 2001; Westerhold et al. 2020).

The medium prevalence of hypoplasia in our sample (∼ 17 %) also suggests good environmental conditions, although some seasonality can be suspected. Indeed, m3 is amongst the most affected loci especially for *P. minutum* (Figure 2, Table 4), and hypoplasia on third molars has been associated with environmental stresses such as seasonality (Franz-Odendaal et al. 2003; Skinner and Pruetz 2012; Upex and Dobney 2012). Periodic floodings are supposed at Ulm-Westtangente and proposed as attritional causes (Heizmann et al. 1989). Such events might result in increased levels of stress through vegetation damage, habitat loss or displacement and decrease of water quality (e.g., Lake et al. 2006), and have already been linked to increased hypoplasia prevalence in rhinocerotids (Hullot et al. 2021). Besides external stressors, an effect of the diet and/or of phylogeny on stress susceptibility has been hypothesized in Miocene rhinocerotids from other sites (Hullot et al. 2023a, b). Here, both species are closely phylogenetically related (early diverging taxa of Aceratheriinae *sensu lato*; Rhinocerotinae *incertae sedis* in Tissier et al. 2020) and are similarly affected despite having different inferred diets (Table 6), suggesting that phylogeny might be the main driver.

## Conclusion

In this article, we provided paleoecological insights for the two rhinocerotid species from the early Miocene locality of Ulm-Westtangente (Germany). All dietary proxies (mesowear, microwear, δ^13^C) pointed towards different feeding preferences, with a more generalistic behavior (mixed-feeding) for the bigger species *Mesaceratherium paulhiacense*. As the rhinocerotids were by far the biggest species at Ulm-Westtangente, competitive interactions with other herbivores might have been limited. However, the investigation of ecological preferences of the other species associated, especially the chalicothere cf. *Metaschizotherium wetzleri* and the tapir *Paratapirus intermedius*, would be interesting to clarify resources use. The prevalence of hypoplasia was similar in both species (∼ 17 %) – suggesting a potential greater influence of phylogeny than diet/ecology in stress susceptibility – but we noted specific differences in the loci affected. Vulnerability periods correlating with life events (birth, juvenile diseases, weaning and cow calf separation, seasonality) were identified in the mortality curves, in accordance with some interpretations on hypoplasia origin. Regarding the paleoenvironment, our rhinocerotid sample gave a mean annual temperature (MAT) of 15.8°C and mean annual precipitation of 317 mm/year suggesting rather warm and dry conditions. This was in agreement with the previous inferences of a warm-temperate forest with grassland patches, but not with the cenogram at the locality, which suggested significantly lower MAT (7°C). Thus, the inclusion of isotopic data from other taxa and/or the use of serial sampling might provide a more robust calculation and an insight into a potential seasonality.

## Acknowledgments

We are grateful to Dr. Eli Amson the curator for fossil mammals at the SMNS for granting access to and providing inventory numbers for the specimens of Ulm-Westtangente. We are also indebted to Jérôme Surault (PALEVOPRIM Poitiers) for scanning several specimens of *P. minutum* used for the microwear analyses.

