## Supplementary 2 for "Life in a Central European warm-temperate to subtropical open forest: paleoecology of the rhinocerotids from Ulm-Westtangente (Aquitanian, early Miocene, Germany)"

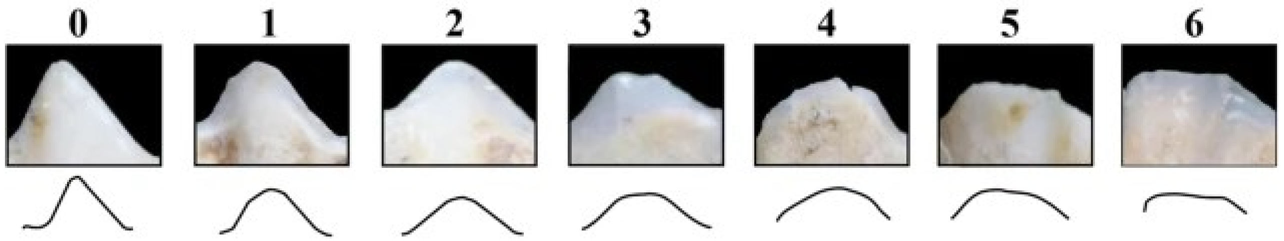


**Figure 1: Mesowear Ruler illustrated with sheep cusps and interpretative drawings**

Adapted from Jiménez-Manchón et al. (2021).

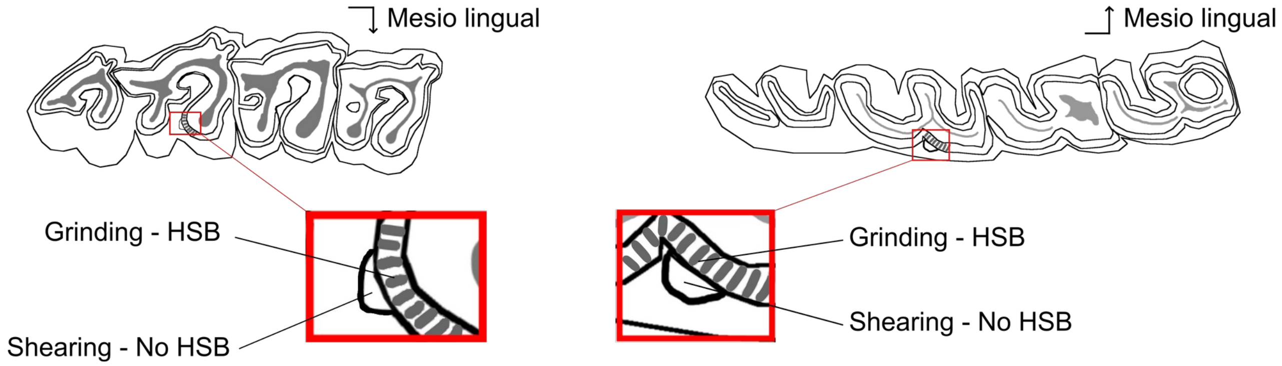


**Figure 2: Localization of the microwear facets on rhinocerotid molars.**

Position of the two microwear facets (grinding and shearing) near the protocone on the second upper molar (left) and near the protoconid on second lower molar (right). Both facets are sampled on the same enamel band with (grinding) or without (shearing) Hunter-Schreger bands (HSB). Modified after Hullot et al. (2019).

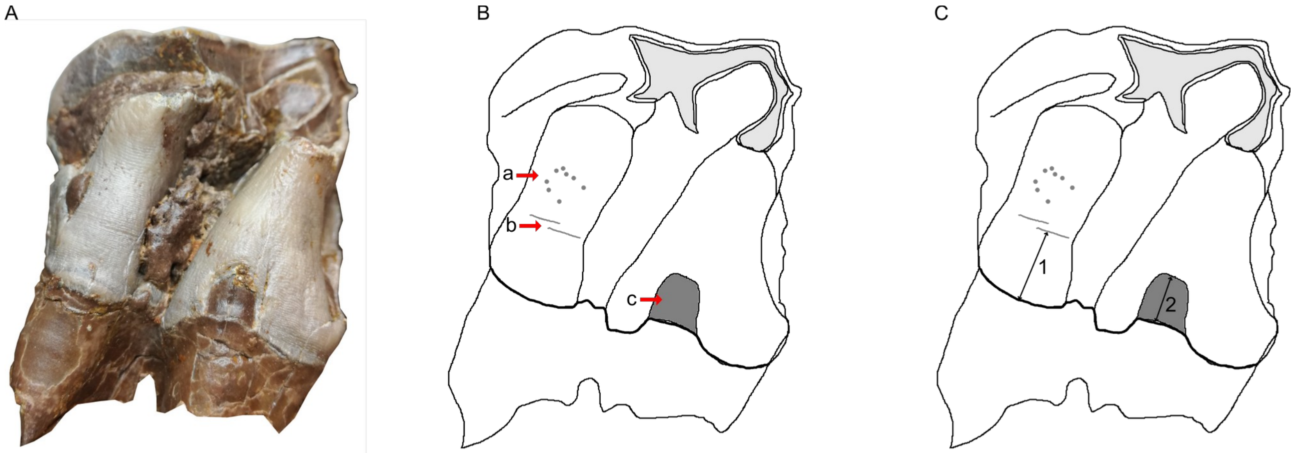


**Figure 3: The three different types of hypoplasia considered in this study and the associated measurements.**

A- Lingual view of right M2 of the specimen MHNT.PAL.2004.0.58 (H. beonense) displaying three types of hypoplasia. B-Interpretative drawing of the photo in A illustrating the hypoplastic defects: a- pitted hypoplasia, b- linear enamel hypoplasia, and c- aplasia. C- Interpretative drawing of the photo in A illustrating the measurements: 1- distance between the base of the defect and the enamel-dentin junction, 2- width of the defect (when applicable).

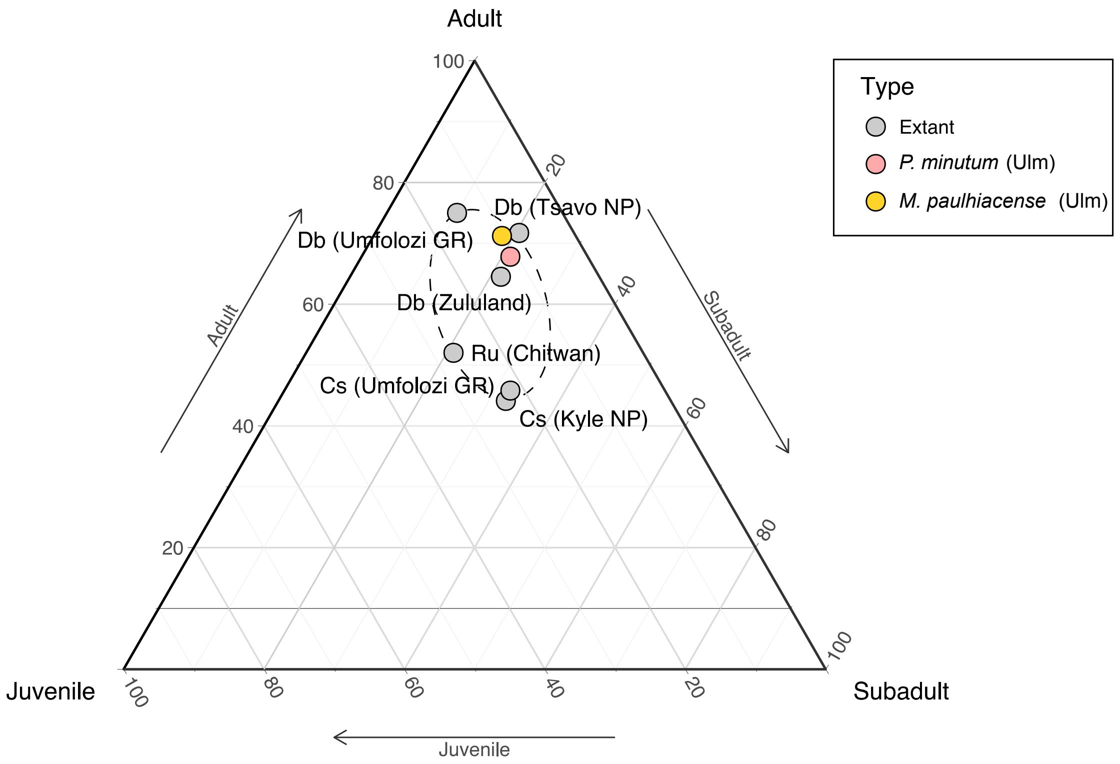


**Figure 4: Percentages of juveniles, subadults and adults in the rhinocerotid samples from Ulm and in various extant populations**
Extant species: Cs – *Ceratotherium simum*, Db – *Diceros bicornis*, Ru – *Rhinoceros unicornis*
References: Chitwan (Laurie et al., 1983), Tsavo National Park (Goddard, 1970), Zululand (Hitchins, 1978), Kyle National Park (Pienaar, 1994), Umfolozi Game Reserve (Pienaar, 1994)

Life curves (qx) of the two species are not different according to a two-sided Kolmogorov-Smirnov test ($D = 0.375$, p-value = 0.2145)

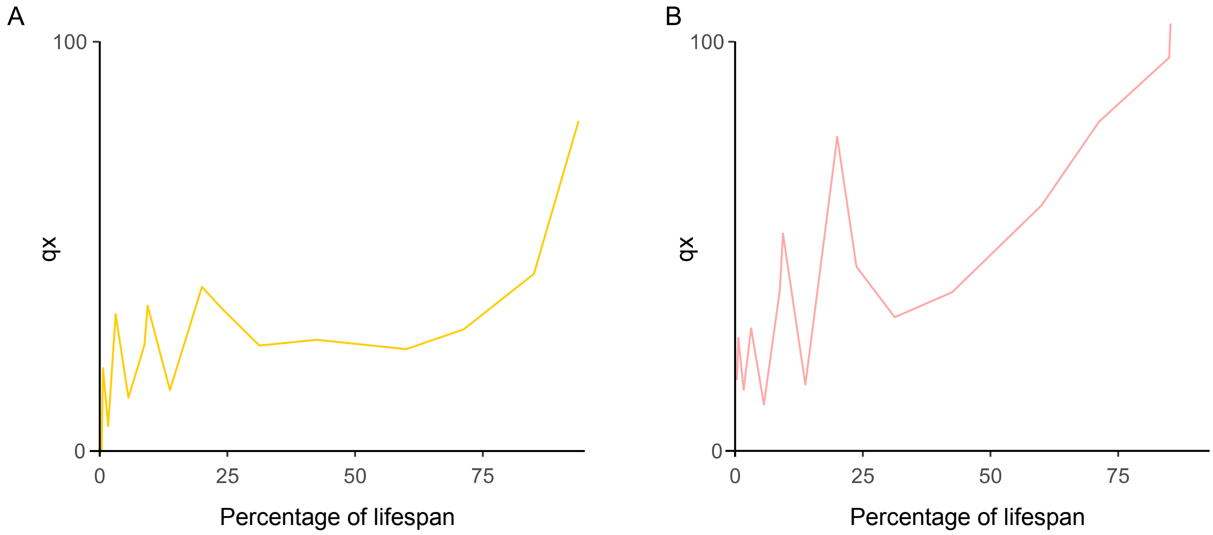


**Figure 5: Mortality rate curves (qx) of the rhinocerotid from Ulm-Westtangente**
qx is the mortality rate calculated as the number of mortalities out of a group of 1000 for each percentage of lifespan: $qx = \frac{dx * 1000}{i * lx}$
where dx = deaths per age class, lx = number of survivors, and i = duration of age class.
A – Mortality rate curve for *M. paulhiacense* from Ulm-Westtangente
B – Mortality rate curve for *P. minutum* from Ulm-Westtangente
