## Supplementary 3 for "Life in a Central European warm-temperate to subtropical open forest: paleoecology of the rhinocerotids from Ulm-Westtangente (Aquitanian, early Miocene, Germany)"

### Ulm-Westangente (SNMS, Stuttgart)

PALEVOPRIM - October 2022 - Manon HULLOT
PALEVOPRIM - October 2023 - Jérôme SURAULT (acquisition), Manon HULLOT (treatment)

***Mesaceratherium paulhiacense***

SMNS-47926
SMNS-47947
SMNS-47954
SMNS-47978
SMNS-47990
SMNS-48071
SMNS-48098
SMNS-48174
SMNS-48126
SMNS-48108
SMNS-48139

***Protaceratherium minutum***

SMNS-47920
SMNS-47963
SMNS-47986
SMNS-47993
SMNS-48055
SMNS-48068
SMNS-48081
SMNS-48089
SMNS-48095
SMNS-48074

October 2023

SMNS-47871
SMNS-47876
SMNS-47977
SMNS-47980
SMNS-48104
SMNS-48107
SMNS-48128
SMNS-48132
SMNS-48134
SMNS-48164
SMNS-48168
SMNS-48176

PS8-Zinv-SNMS_47926-um3s-ptc-hsb

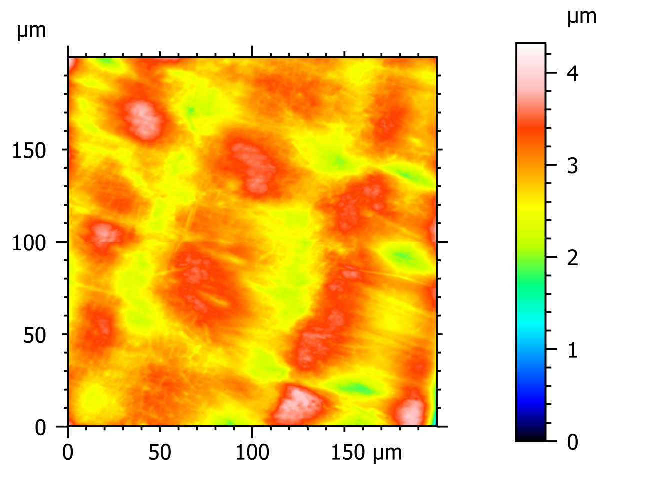


PS8-Zinv-SNMS_47926-um3s-ptc-hsb

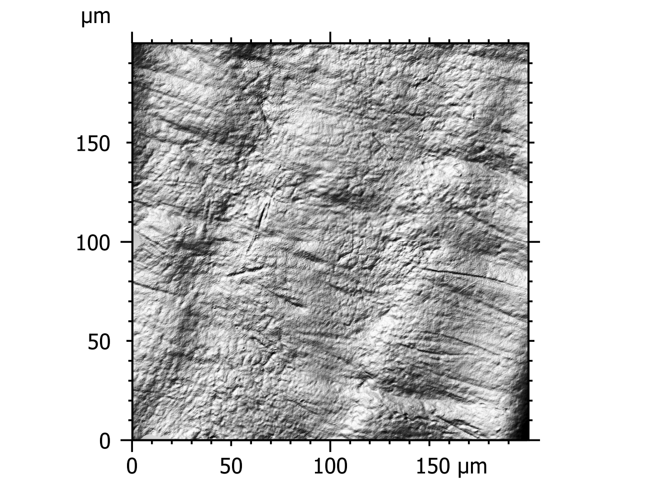


PS8-Zinv-SNMS_47926-um3s-ptc-nohsb

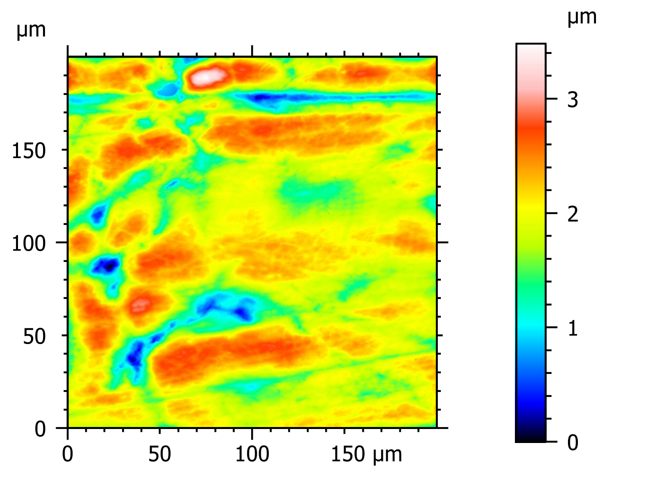


PS8-Zinv-SNMS_47926-um3s-ptc-nohsb

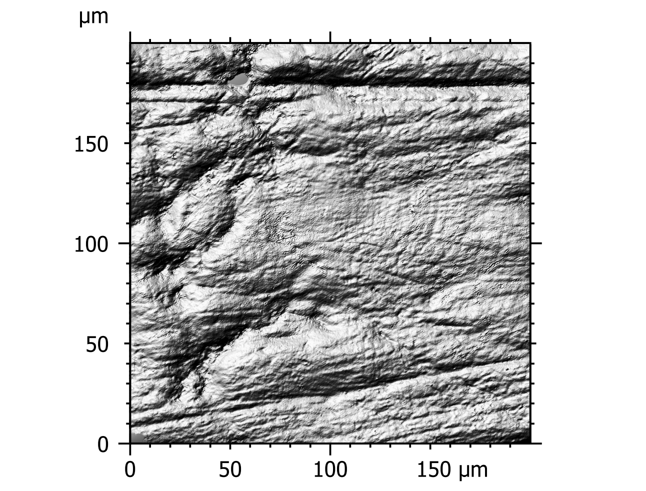


PS8-Zinv-SNMS_47947-um1s-ptc-hsb

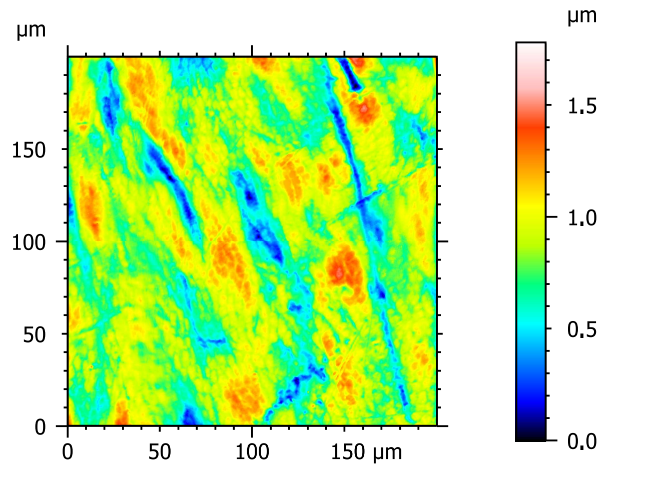


PS8-Zinv-SNMS_47947-um1s-ptc-hsb

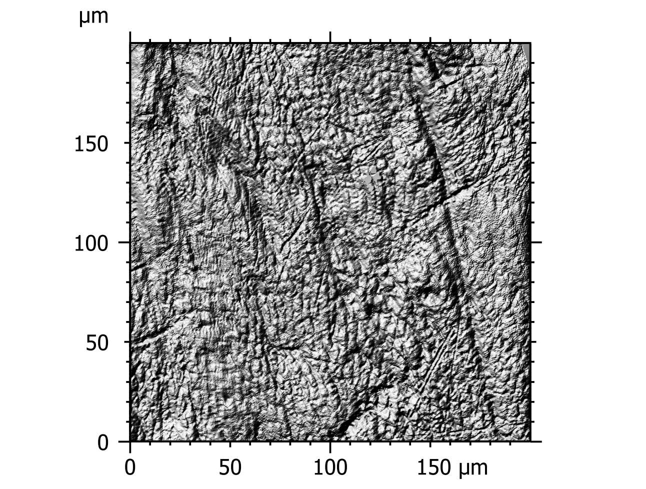

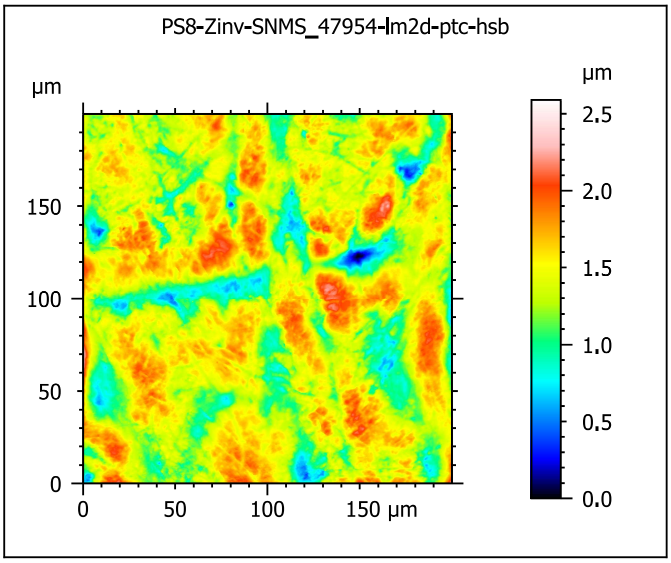


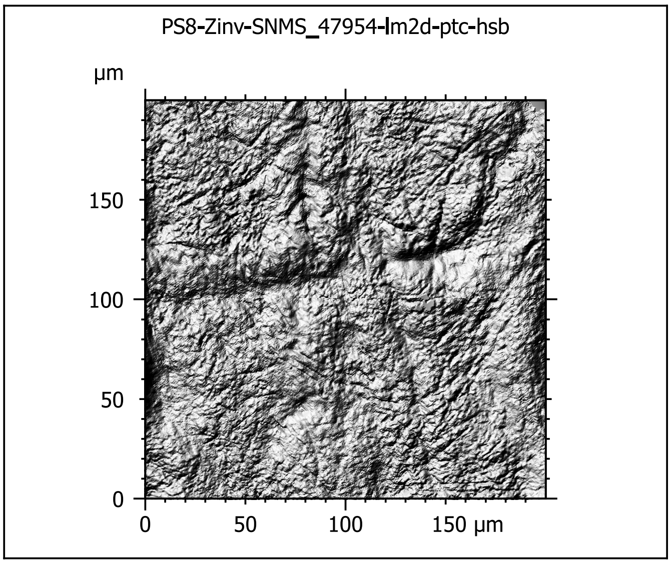


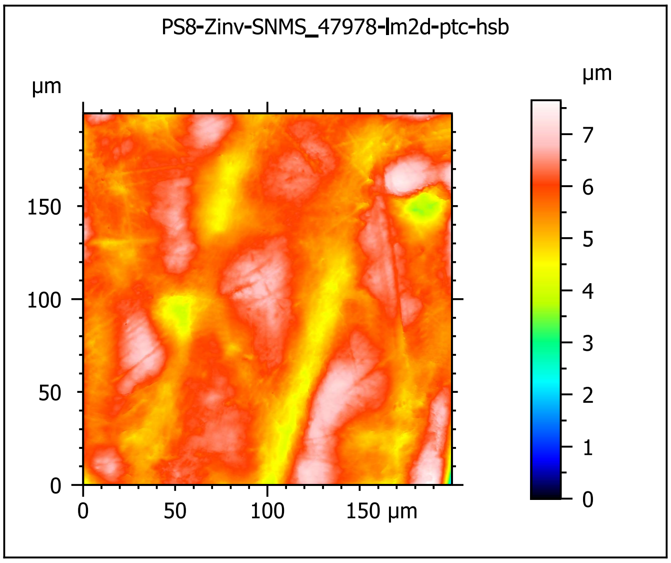


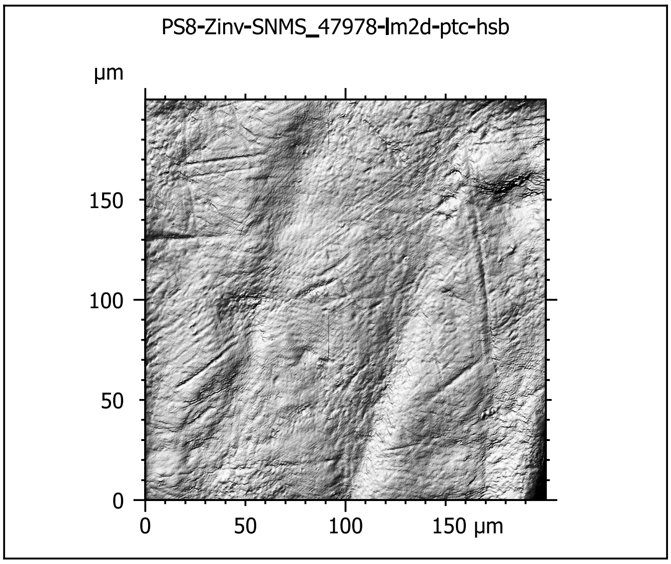


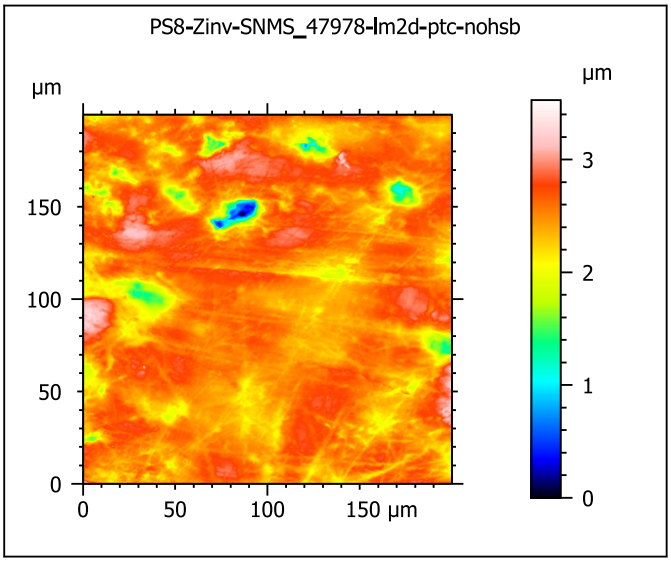


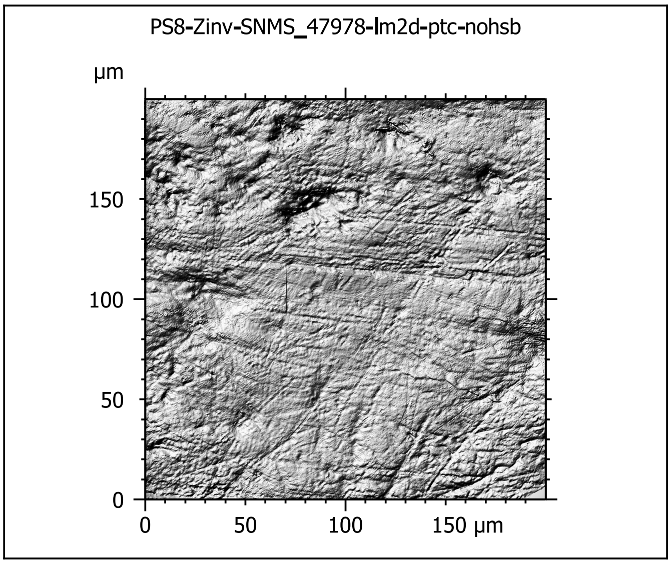


MountainsMap® Premium 10.0.10510

PS8-Zinv-SNMS_47990-lm2d-hyp-nohsb

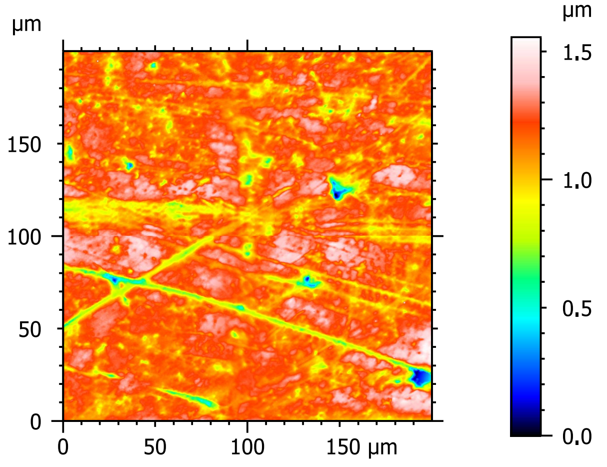


PS8-Zinv-SNMS_47990-lm2d-hyp-nohsb

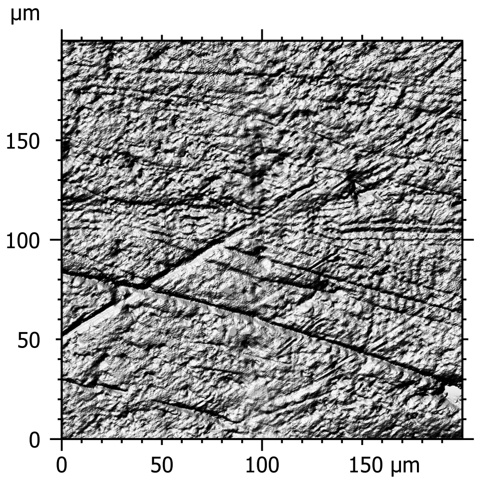


PS8-Zinv-SNMS_47990-lm2d-ptcd-hsb

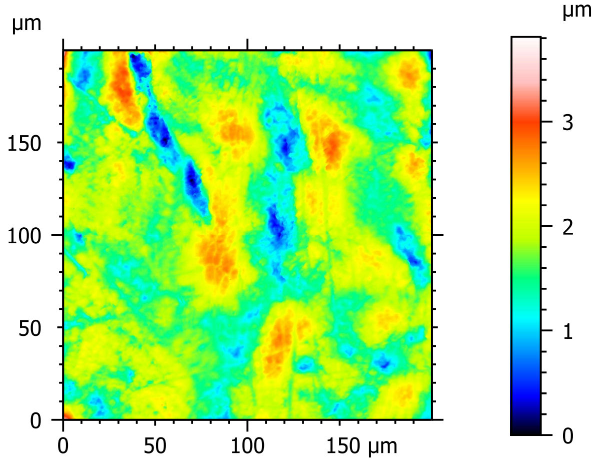


PS8-Zinv-SNMS_47990-lm2d-ptcd-hsb

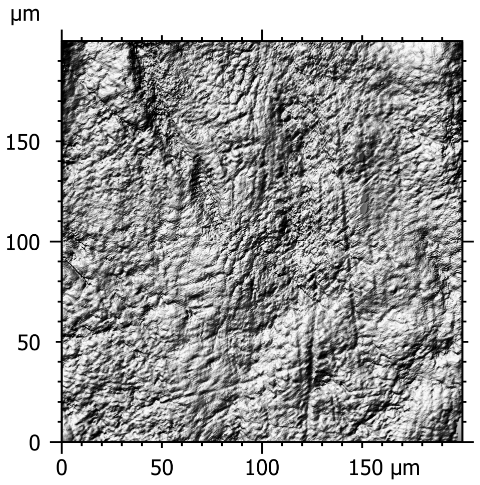


PS8-Zinv-SNMS_48071-lm3d-ptcd-hsb

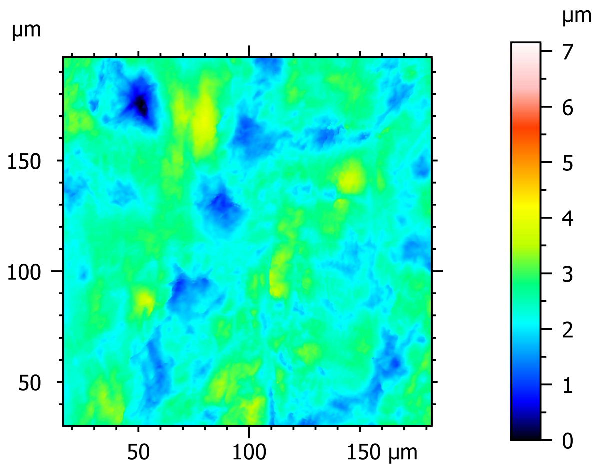


PS8-Zinv-SNMS_48071-lm3d-ptcd-hsb

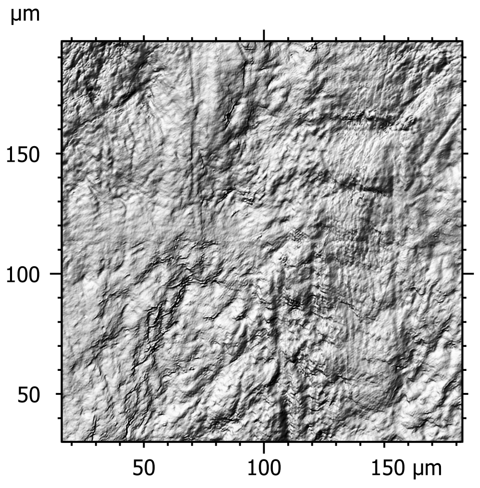


MountainsMap® Premium 10.0.10510

PS8-Zinv-SNMS_48098-um1s-ptc-hsb

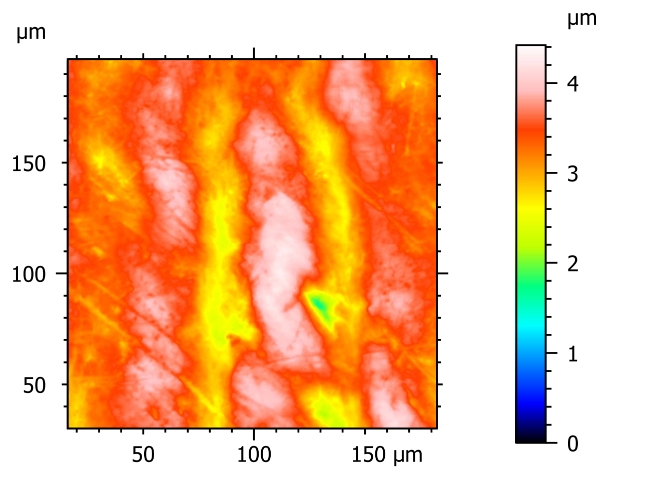


PS8-Zinv-SNMS_48098-um1s-ptc-hsb

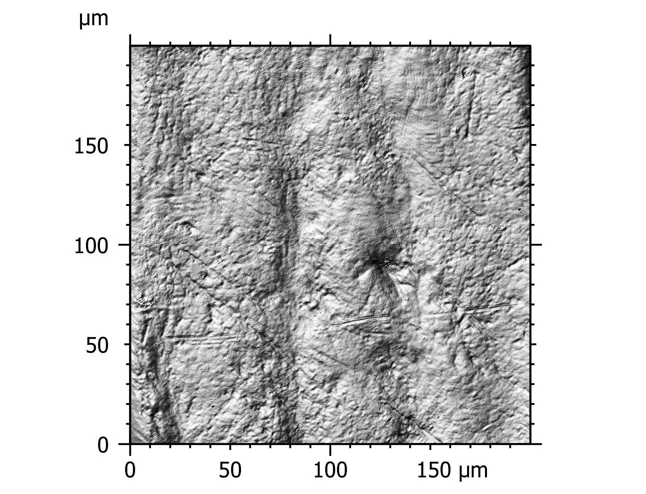


PS8-Zinv-SNMS_48098-um1s-ptc-nohsb

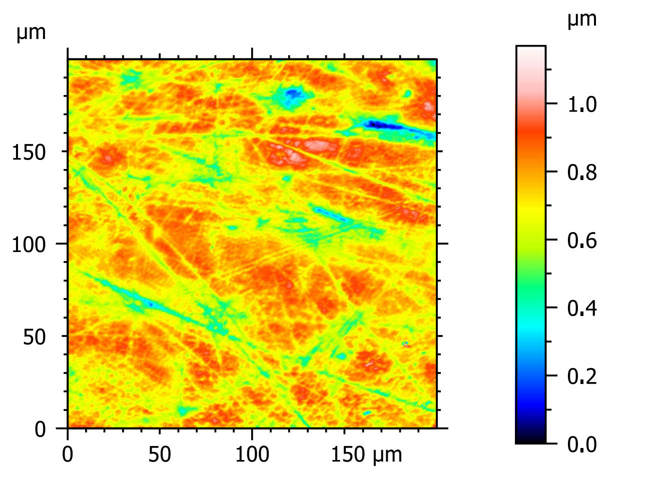


PS8-Zinv-SNMS_48098-um1s-ptc-nohsb

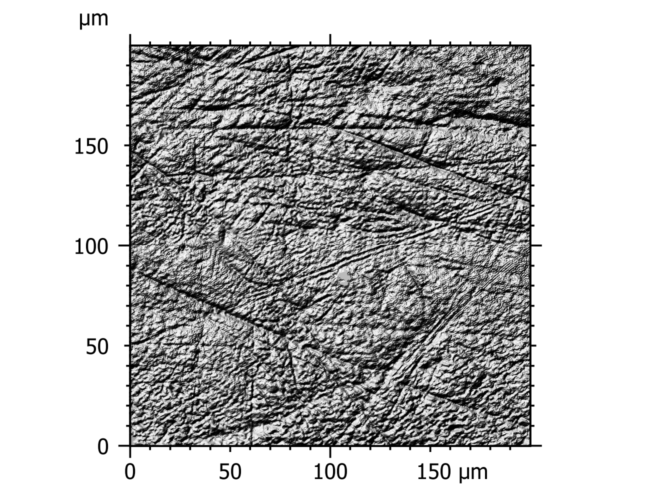


PS8-Zinv-SNMS_48108-um3d-ptc-hsb

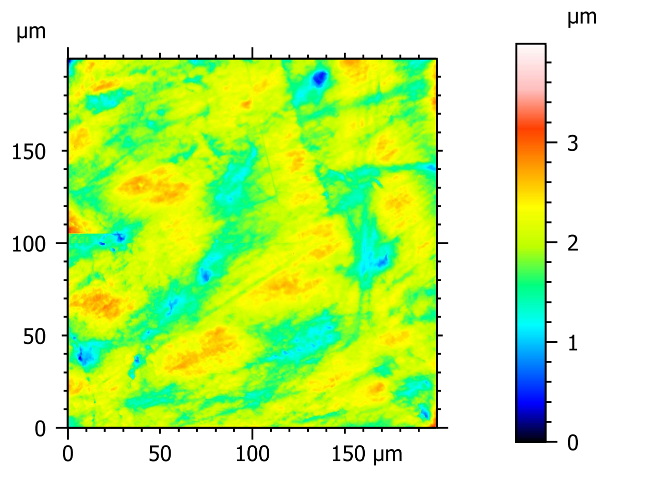


PS8-Zinv-SNMS_48108-um3d-ptc-hsb

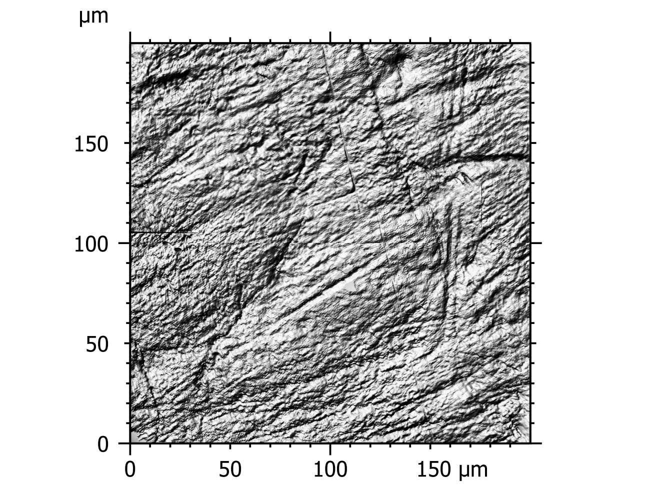


MountainsMap® Premium 10.0.10510

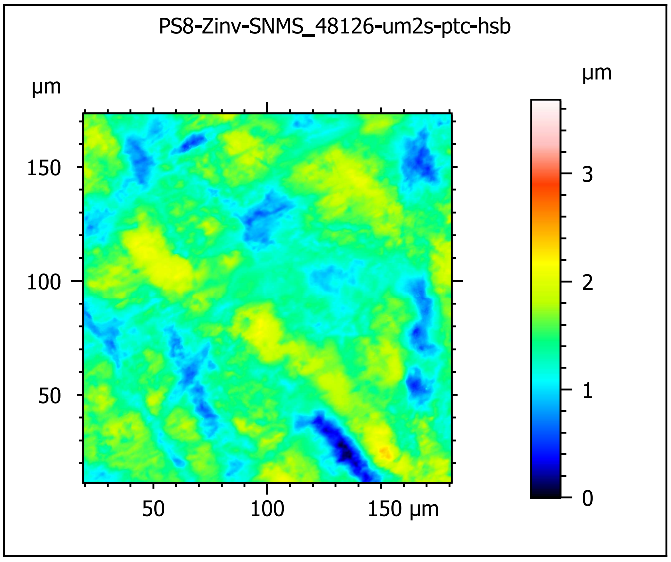
PS8-Zinv-SNMS_48126-um2s-ptc-hsb
µm
150
100
50
50
100
150 µm
µm
3
2
1
0

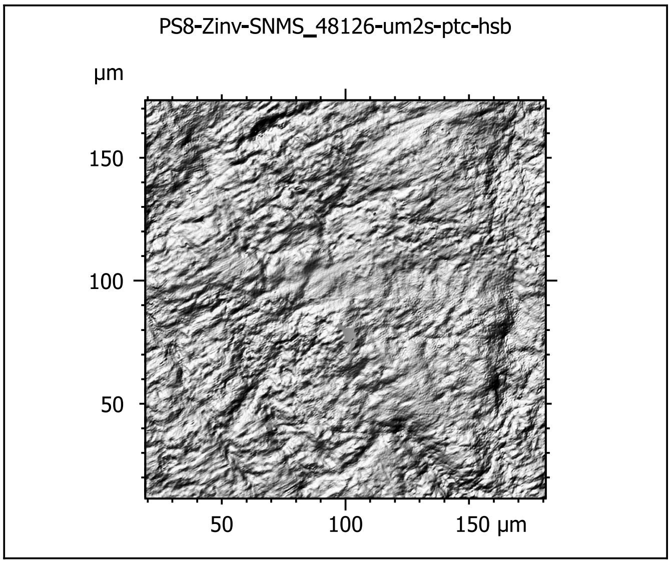
PS8-Zinv-SNMS_48126-um2s-ptc-hsb
µm
150
100
50
50
100
150 µm

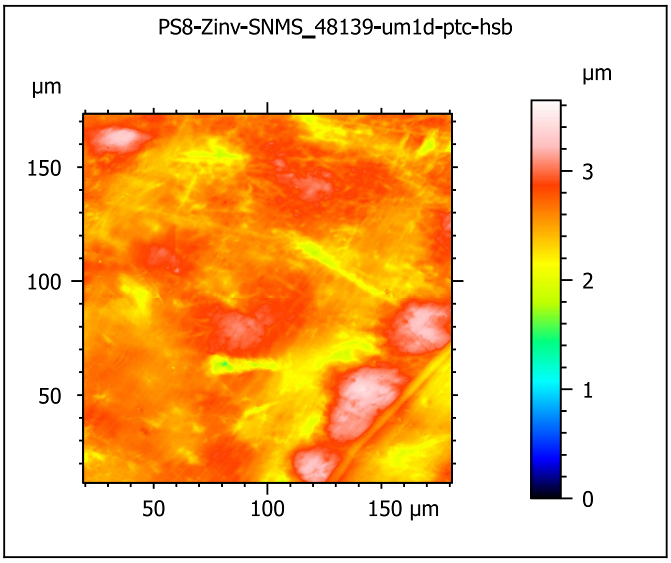
PS8-Zinv-SNMS_48139-um1d-ptc-hsb
µm
150
100
50
50
100
150 µm
µm
3
2
1
0

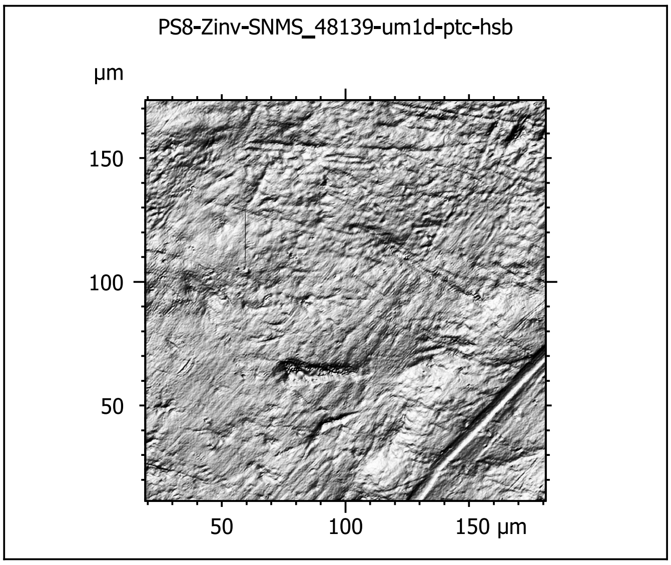
PS8-Zinv-SNMS_48139-um1d-ptc-hsb
µm
150
100
50
50
100
150 µm

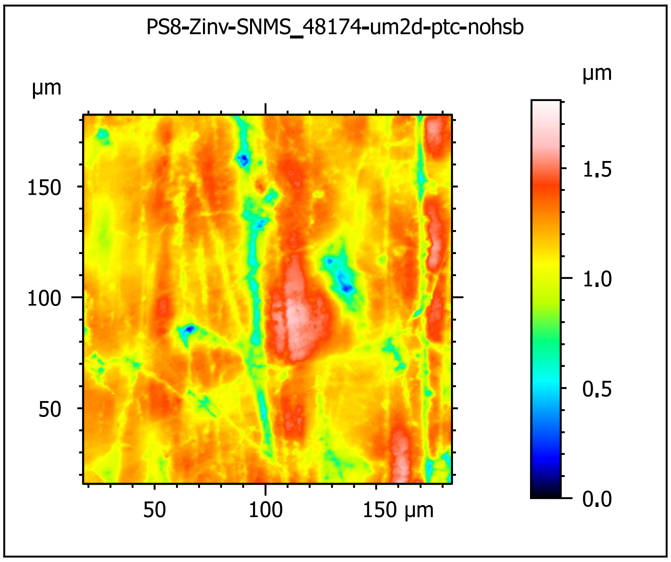
PS8-Zinv-SNMS_48174-um2d-ptc-nohsb
µm
150
100
50
50
100
150 µm
µm
1.5
1.0
0.5
0.0

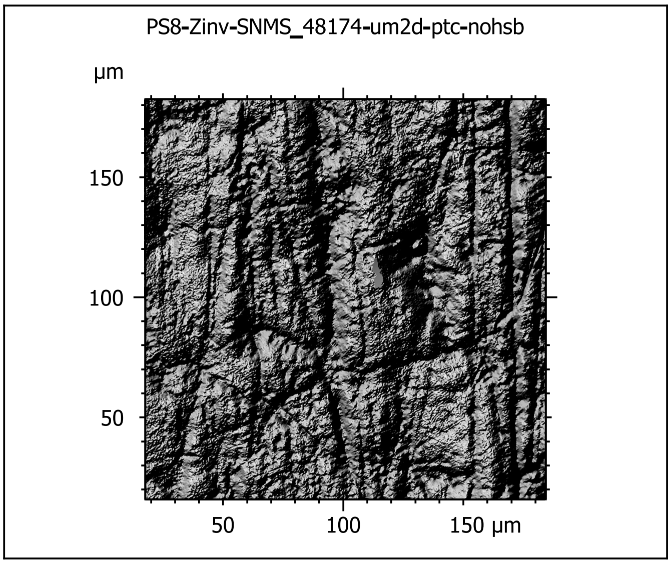
PS8-Zinv-SNMS_48174-um2d-ptc-nohsb
µm
150
100
50
50
100
150 µm

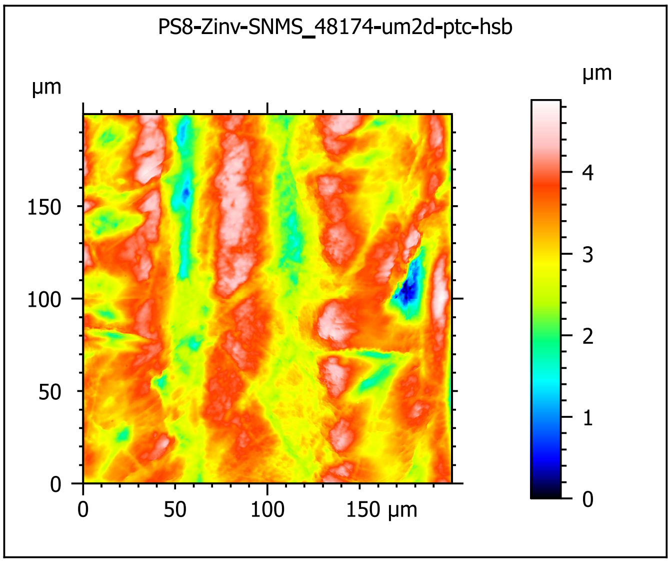
PS8-Zinv-SNMS_48174-um2d-ptc-hsb
µm
150
100
50
0
0
50
100
150 µm
µm
4
3
2
1
0

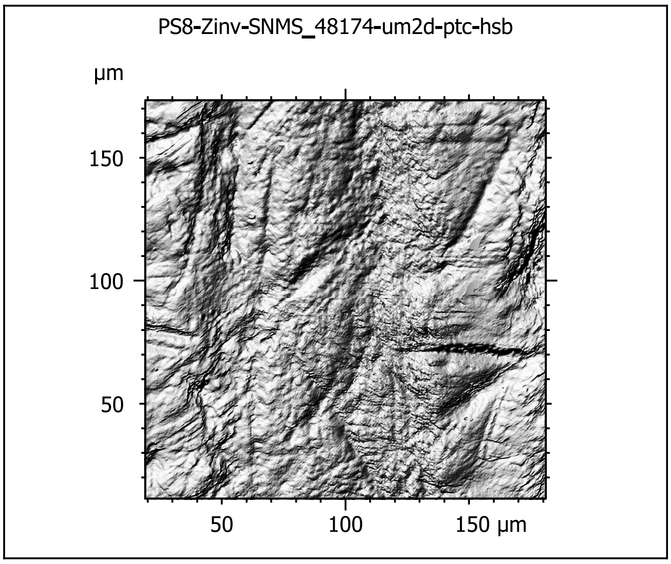
PS8-Zinv-SNMS_48174-um2d-ptc-hsb
µm
150
100
50
50
100
150 µm

PS8-Zinv-SNMS_47920-lm1s-ptcd-hsb

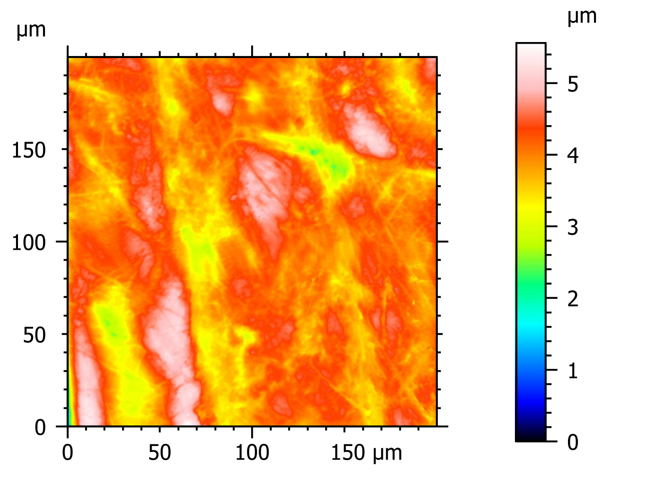


PS8-Zinv-SNMS_47920-lm1s-ptcd-hsb

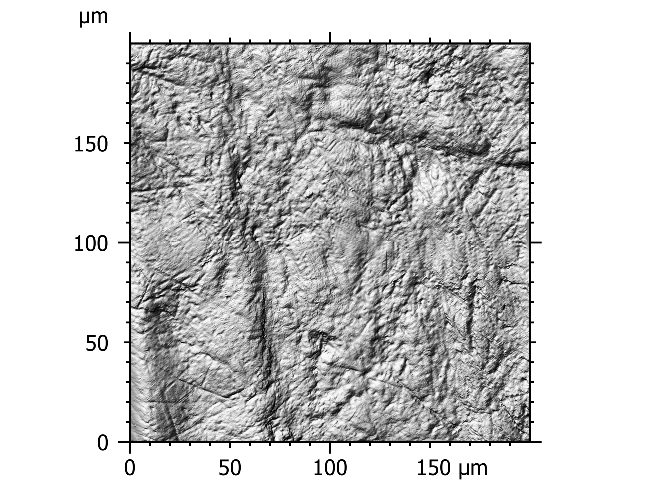


PS8-Zinv-SNMS_47963-um2s-ptc-hsb

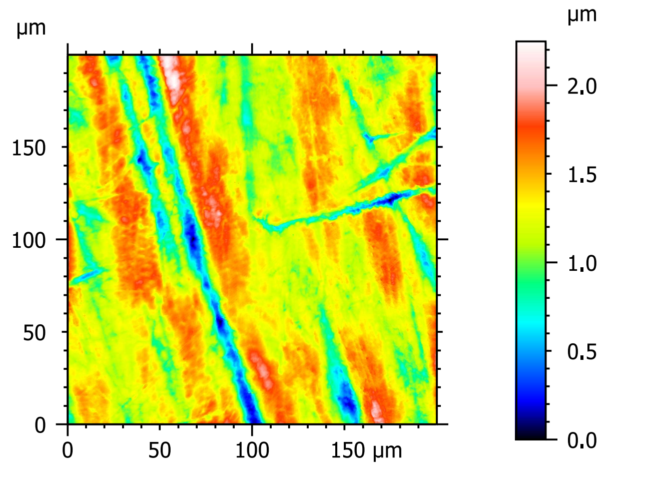


PS8-Zinv-SNMS_47963-um2s-ptc-hsb

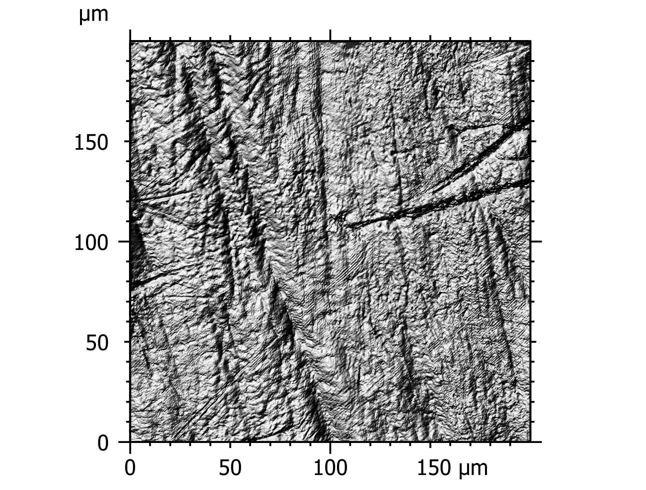


PS8-Zinv-SNMS_47986-lm1d-hyp-hsb

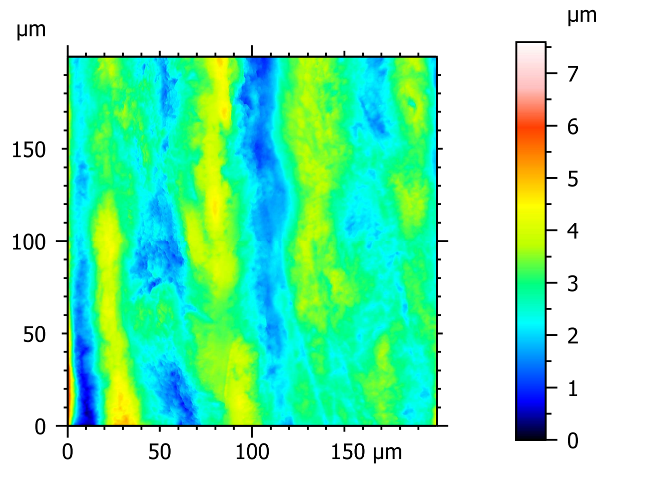


PS8-Zinv-SNMS_47986-lm1d-hyp-hsb

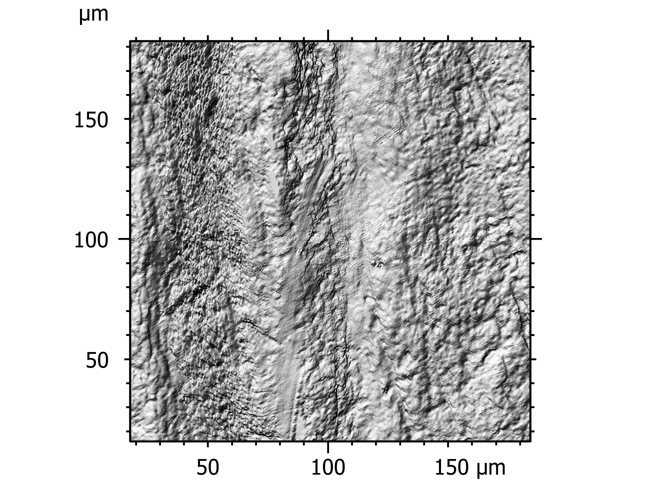


MountainsMap® Premium 10.0.10510

PS8-Zinv-SNMS_47993-um1d-ptc-hsb

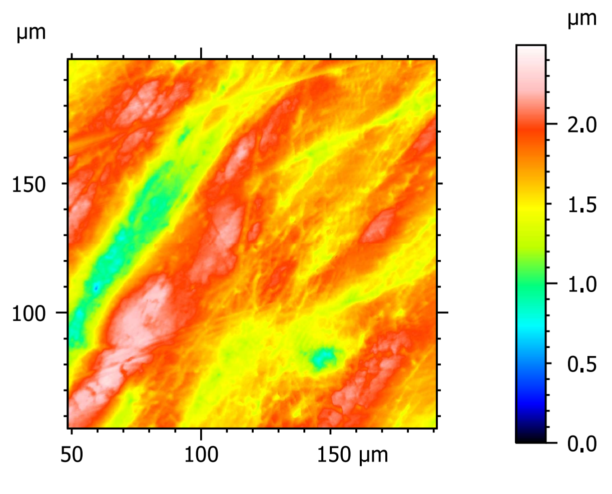


PS8-Zinv-SNMS_47993-um1d-ptc-hsb

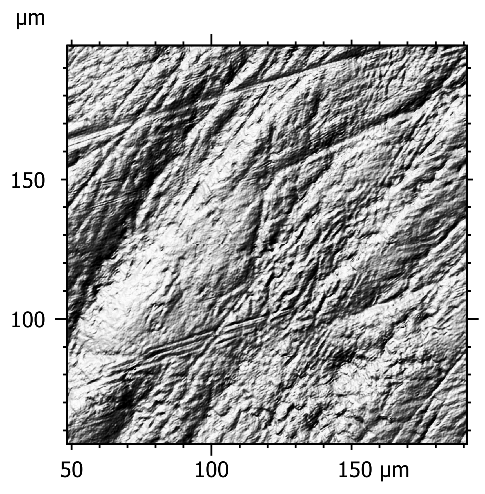


PS8-Zinv-SNMS_48055-lm2s-hyp-hsb

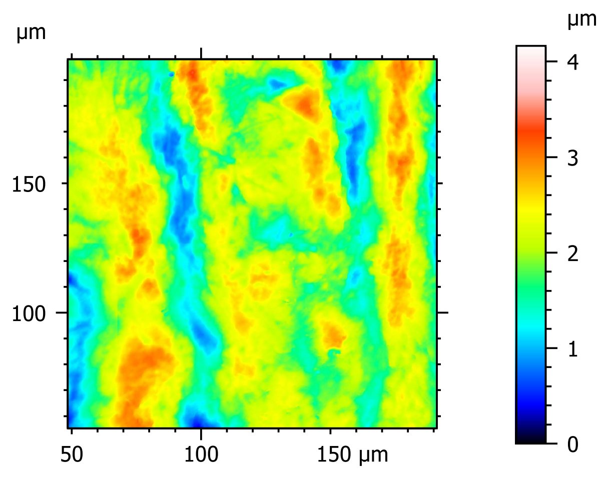


PS8-Zinv-SNMS_48055-lm2s-hyp-hsb

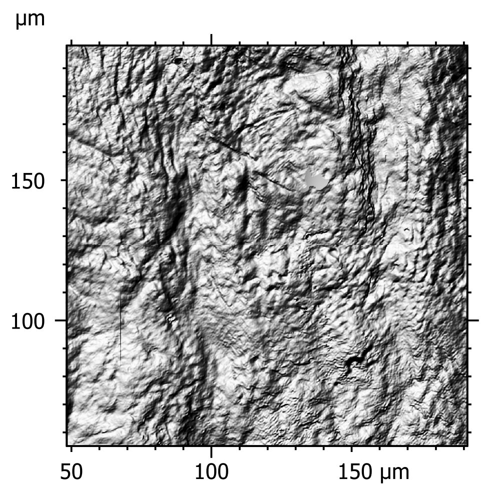


PS8-Zinv-SNMS_48055-um1d-ptc-nohsb

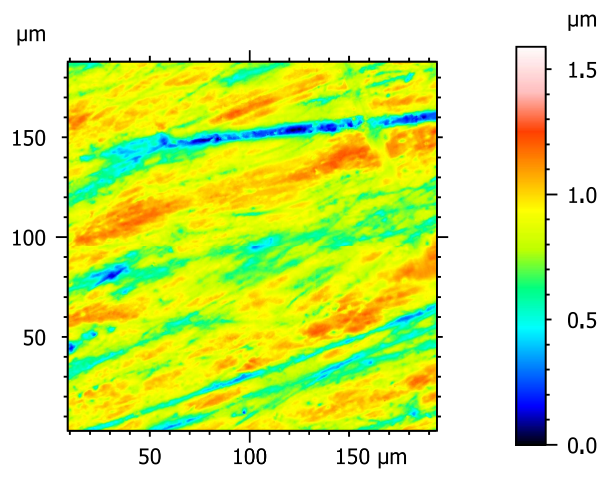


PS8-Zinv-SNMS_48055-um1d-ptc-nohsb

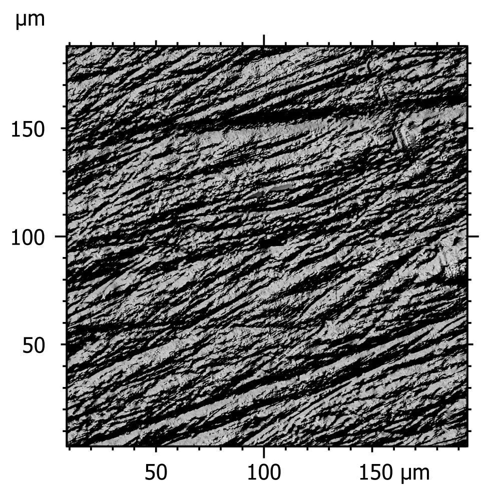


MountainsMap® Premium 10.0.10510

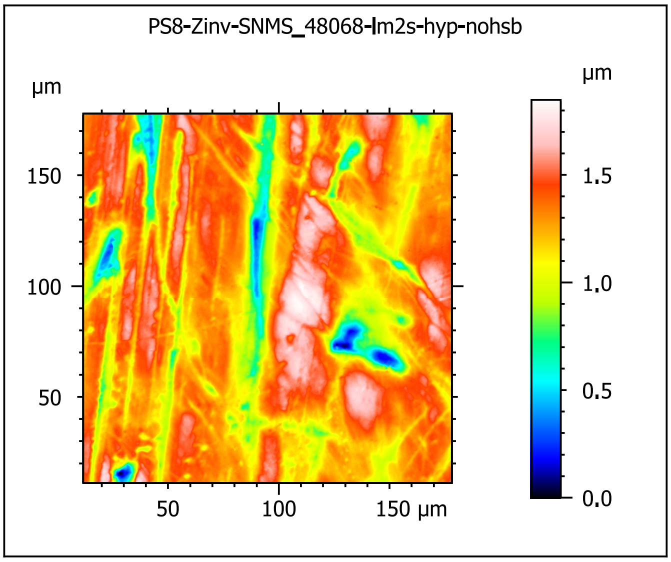


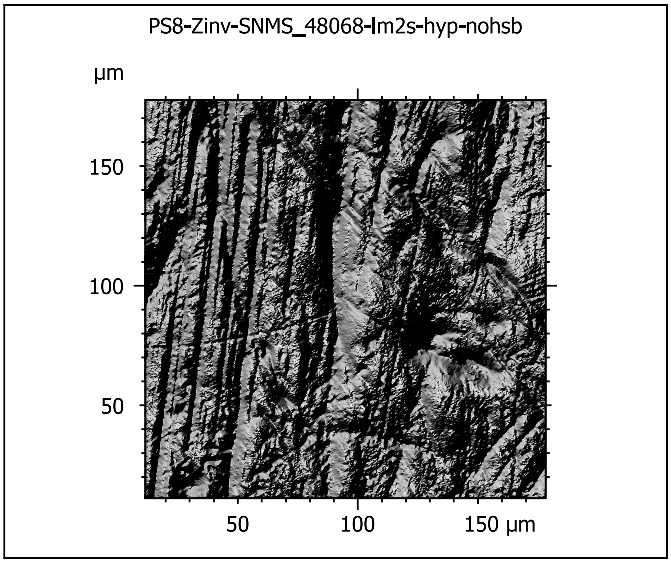


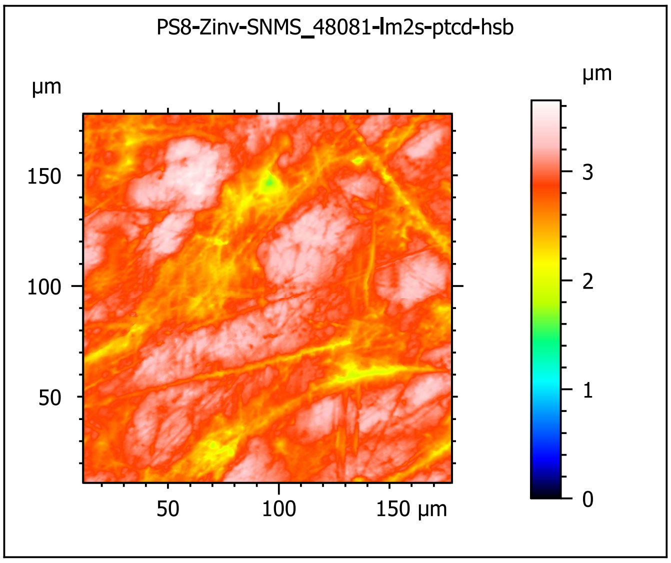


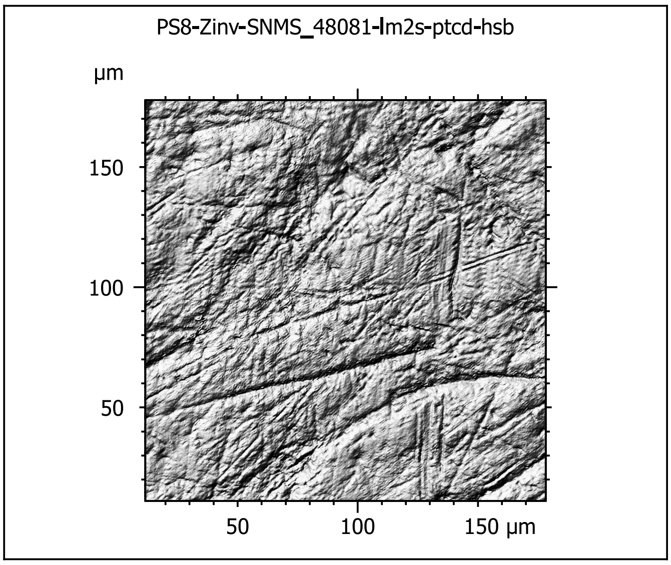


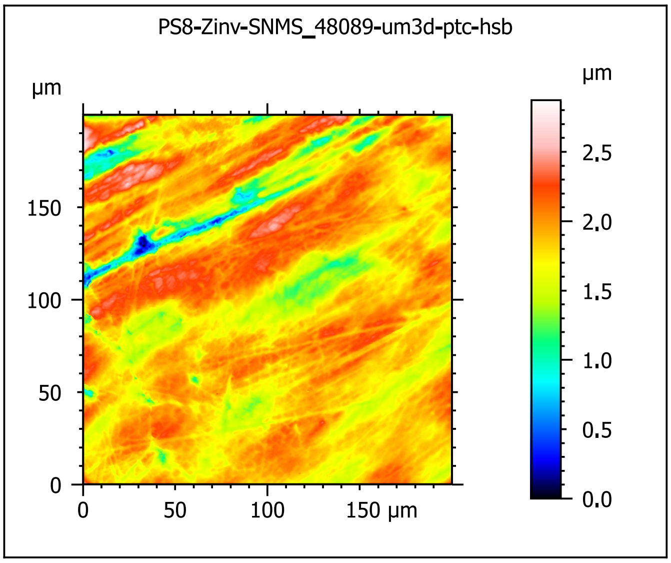


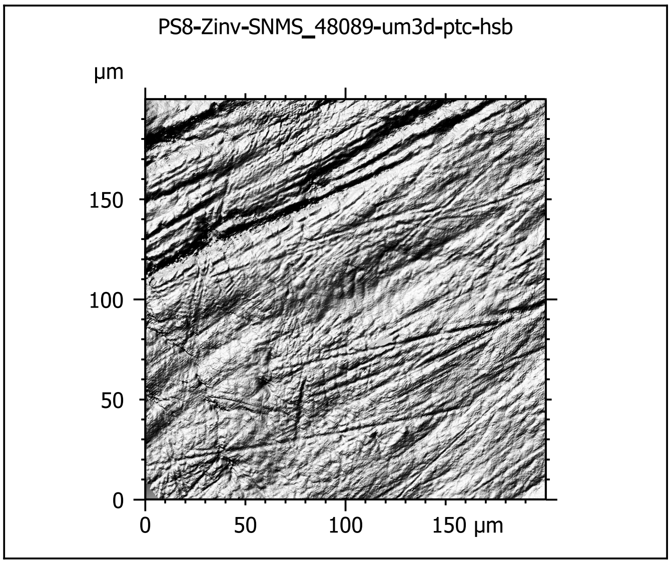


MountainsMap® Premium 10.0.10510

PS8-Zinv-SNMS_48089-um3d-ptc-nohsb

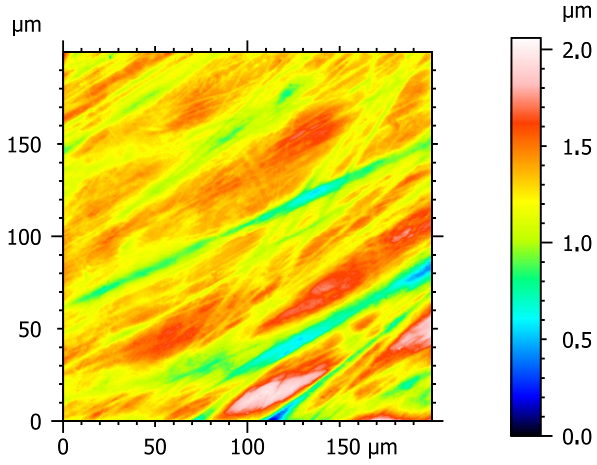


PS8-Zinv-SNMS_48089-um3d-ptc-nohsb

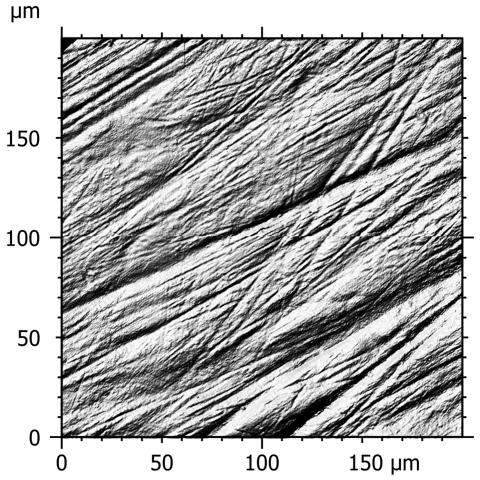


PS8-Zinv-SNMS_48095-lm1s-hyp-hsb

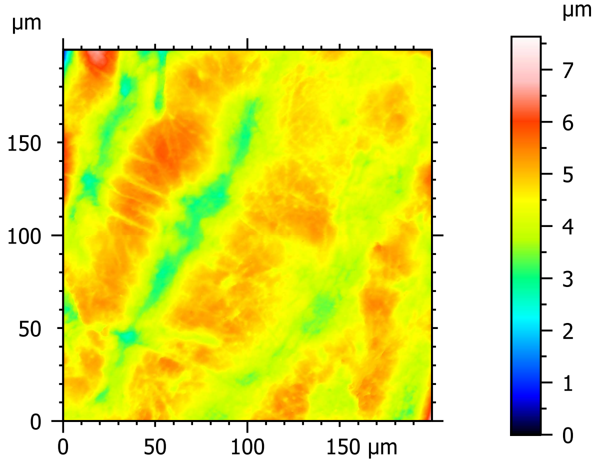


PS8-Zinv-SNMS_48095-lm1s-hyp-hsb

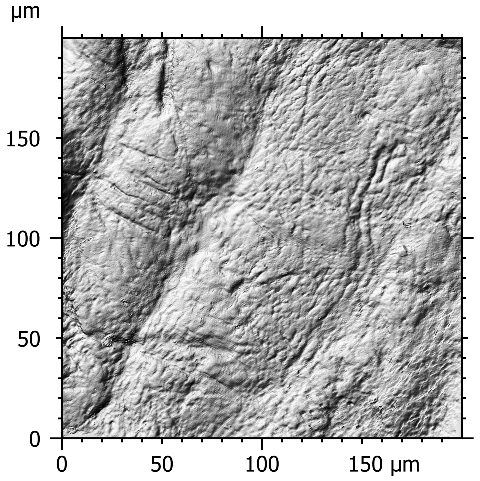


PS8-Zinv-SNMS_48074-lm1s-ptcd-hsb

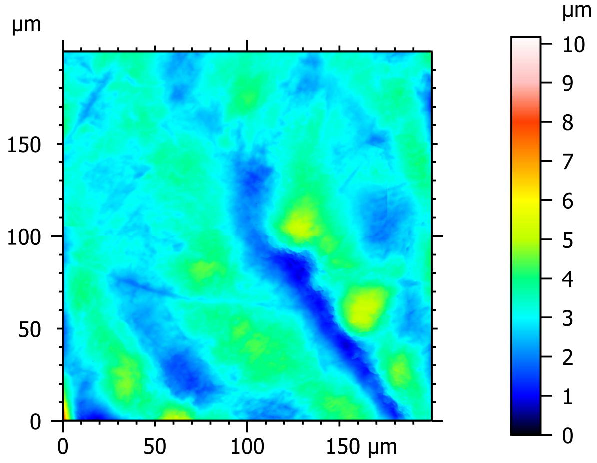


PS8-Zinv-SNMS_48074-lm1s-ptcd-hsb

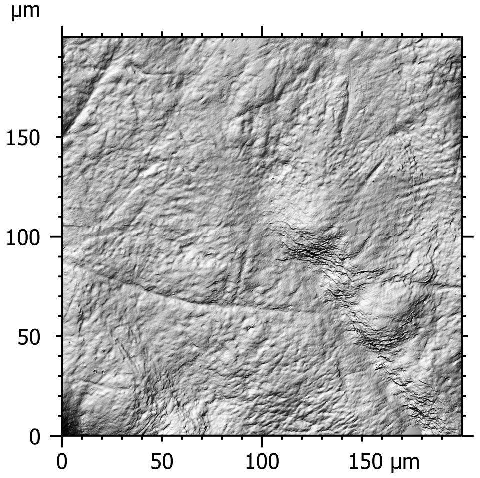


MountainsMap® Premium 10.0.10510

PS8-Zinv-SNMS_47871-UM2d-ptc-hsb

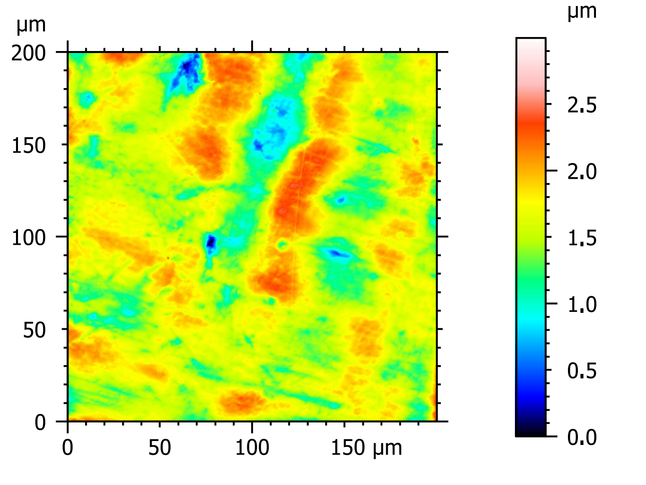


PS8-Zinv-SNMS_47871-UM2d-ptc-hsb

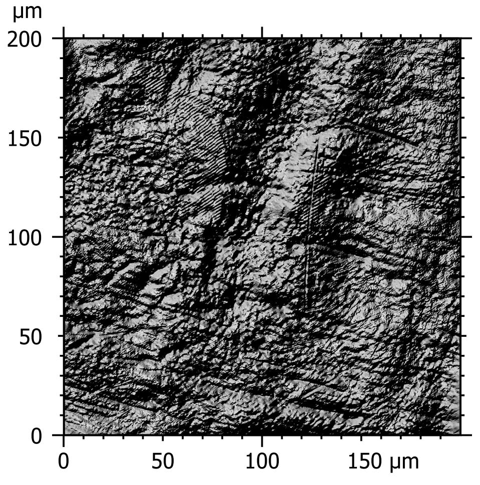


PS8-Zinv-SNMS_47871-UM2s-ptc-no-hsb

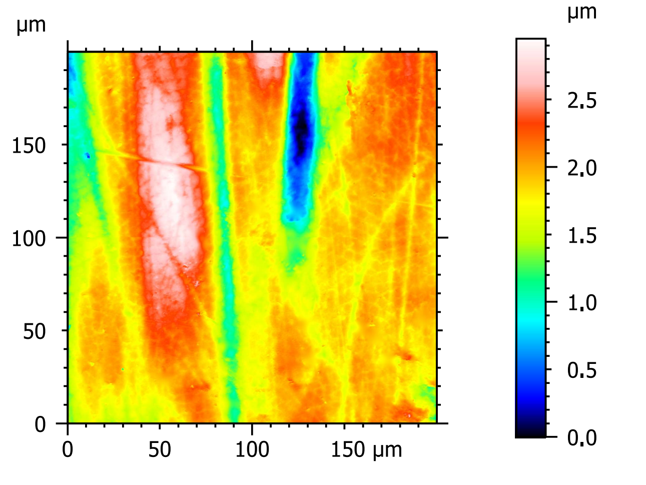


PS8-Zinv-SNMS_47871-UM2s-ptc-no-hsb

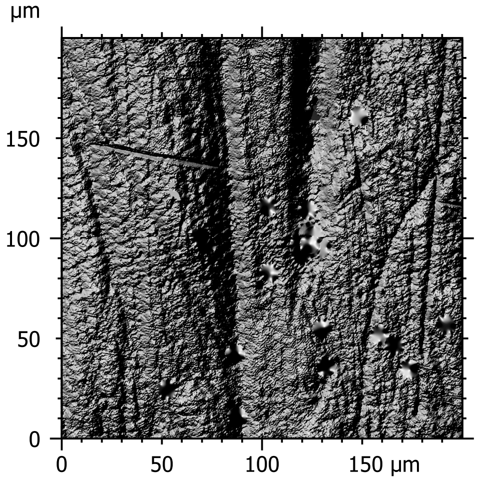


PS8-Zinv-SNMS_47876-UM2d-ptc-hsb

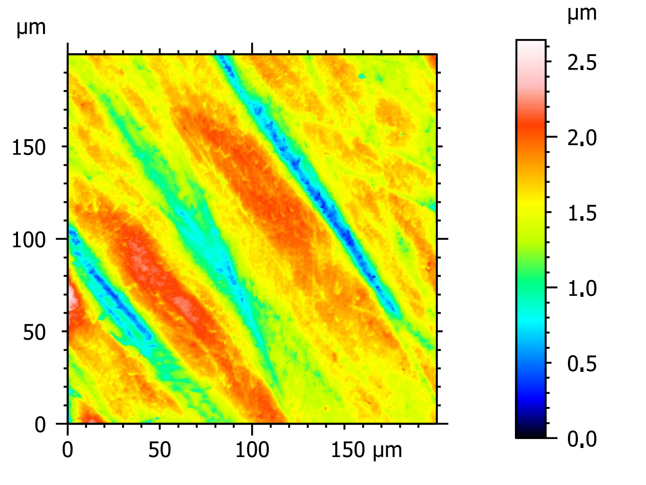


PS8-Zinv-SNMS_47876-UM2d-ptc-hsb

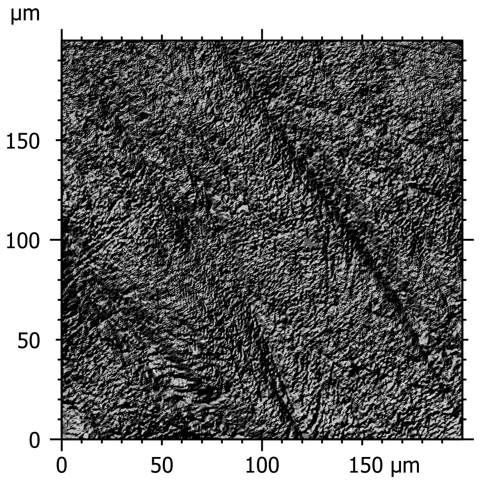

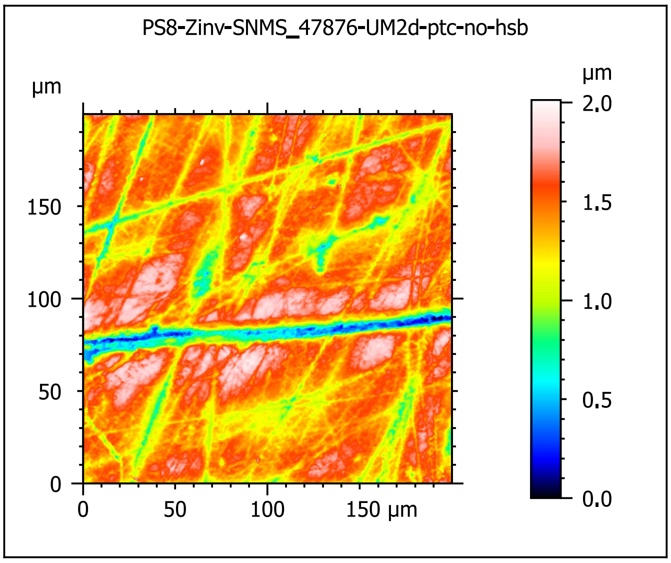


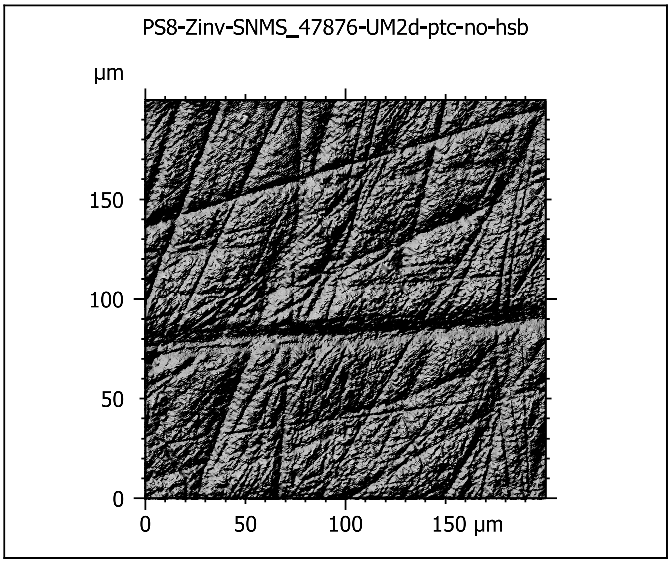


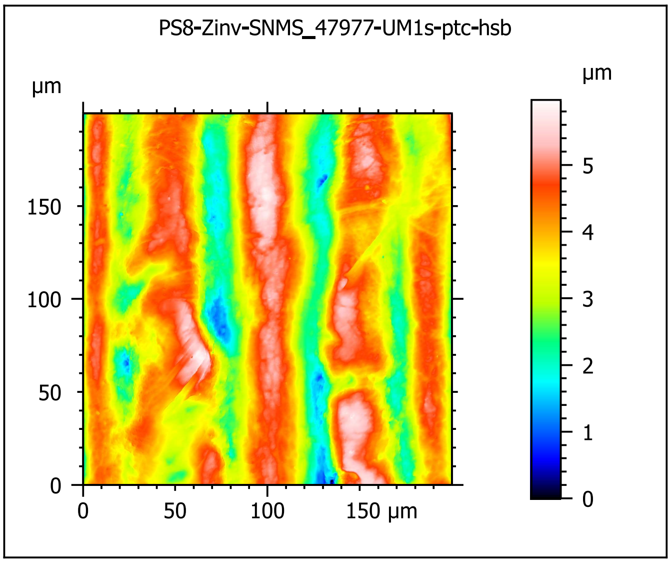


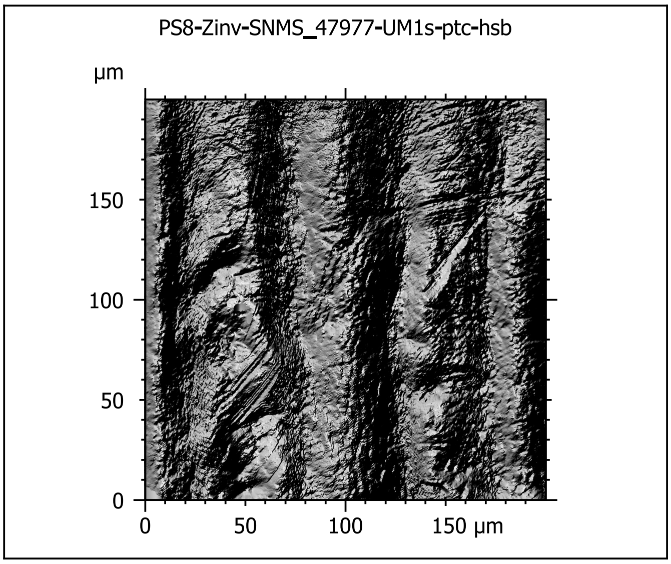


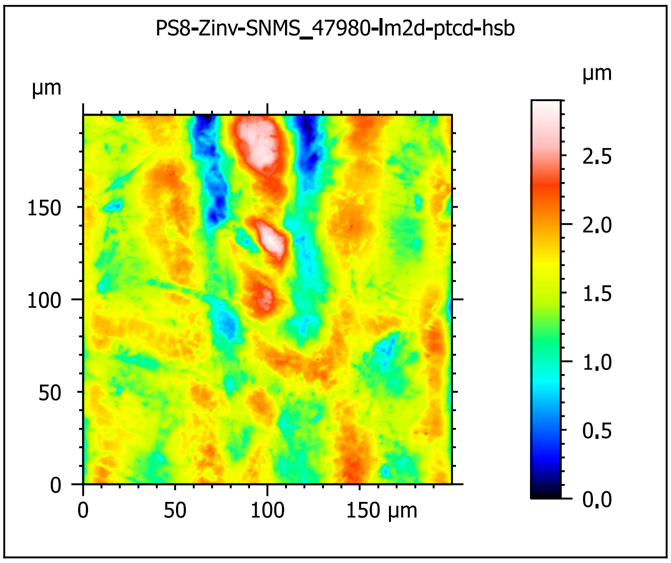


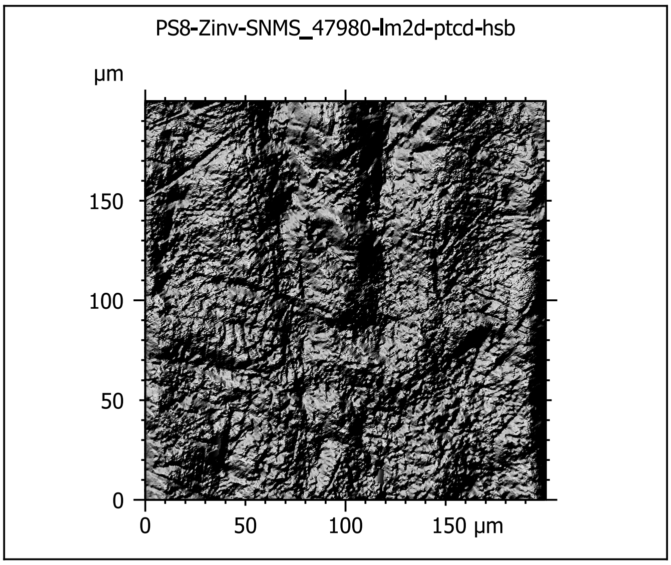


MountainsMap® Premium 10.0.10510

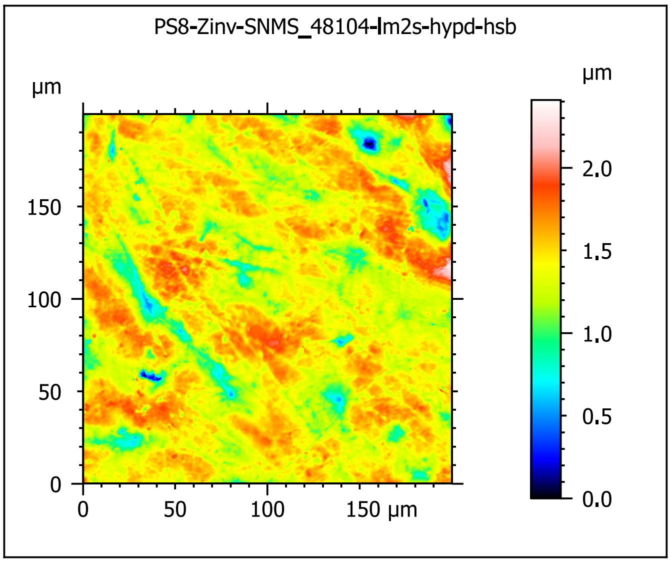
PS8-Zinv-SNMS_48104-lm2s-hypd-hsb
µm
150
100
50
0
0
50
100
150 µm
µm
2,0
1,5
1,0
0,5
0,0

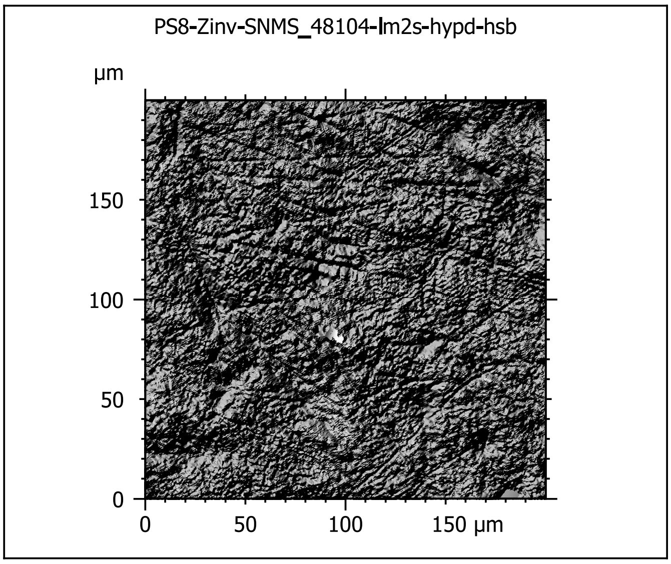
PS8-Zinv-SNMS_48104-lm2s-hypd-hsb
µm
150
100
50
0
0
50
100
150 µm

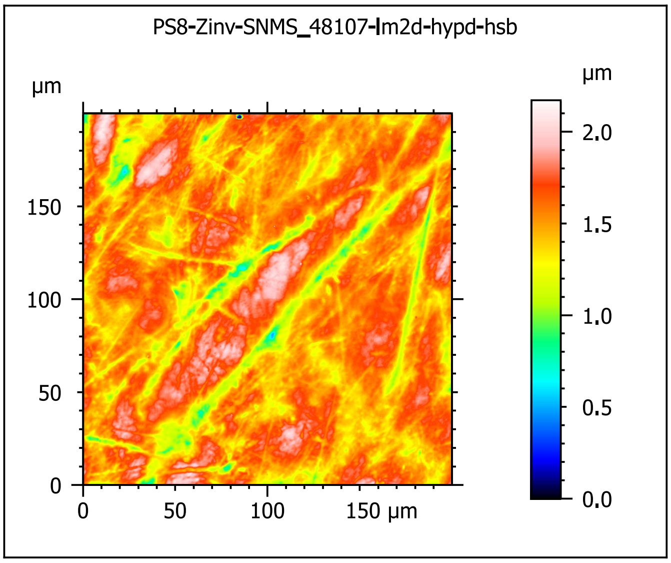
PS8-Zinv-SNMS_48107-lm2d-hypd-hsb
µm
150
100
50
0
0
50
100
150 µm
µm
2,0
1,5
1,0
0,5
0,0

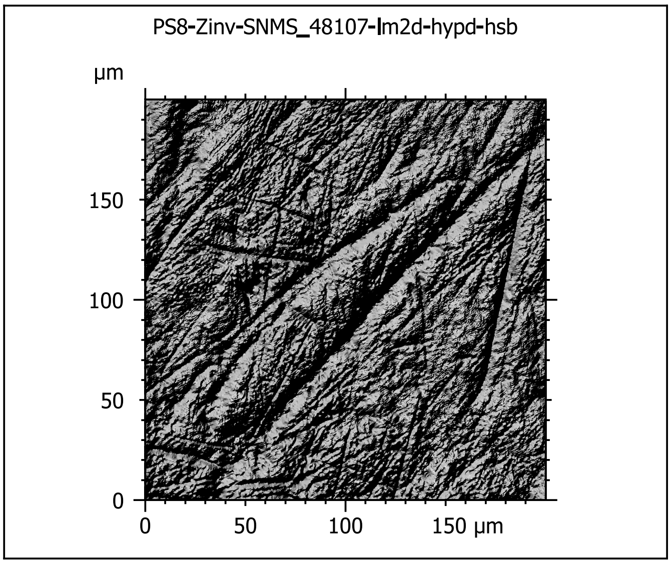
PS8-Zinv-SNMS_48107-lm2d-hypd-hsb
µm
150
100
50
0
0
50
100
150 µm

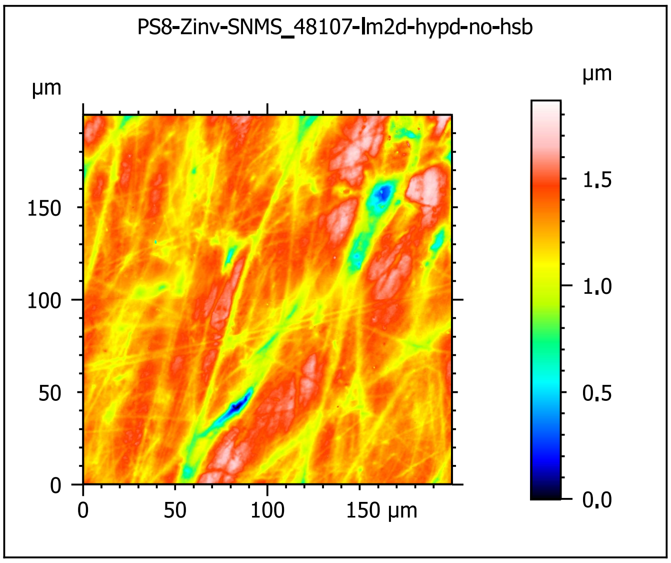
PS8-Zinv-SNMS_48107-lm2d-hypd-no-hsb
µm
150
100
50
0
0
50
100
150 µm
µm
1,5
1,0
0,5
0,0

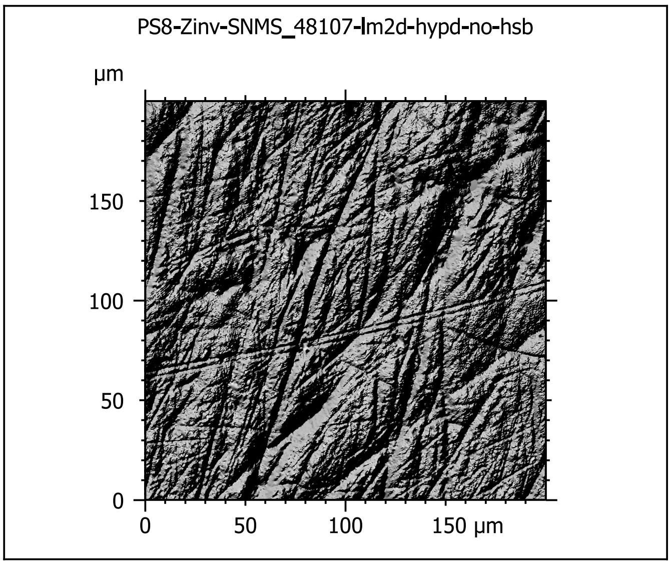
PS8-Zinv-SNMS_48107-lm2d-hypd-no-hsb
µm
150
100
50
0
0
50
100
150 µm

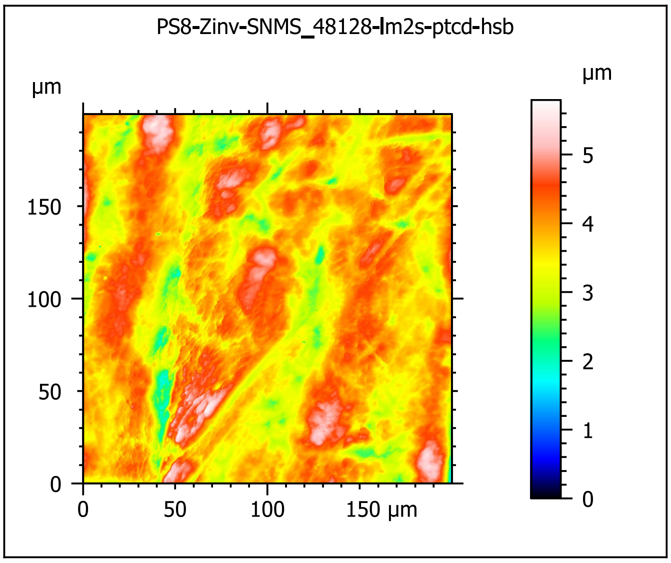


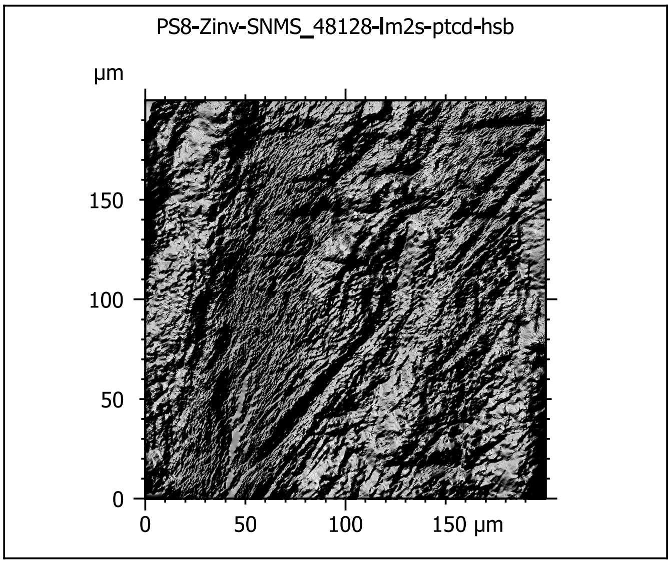


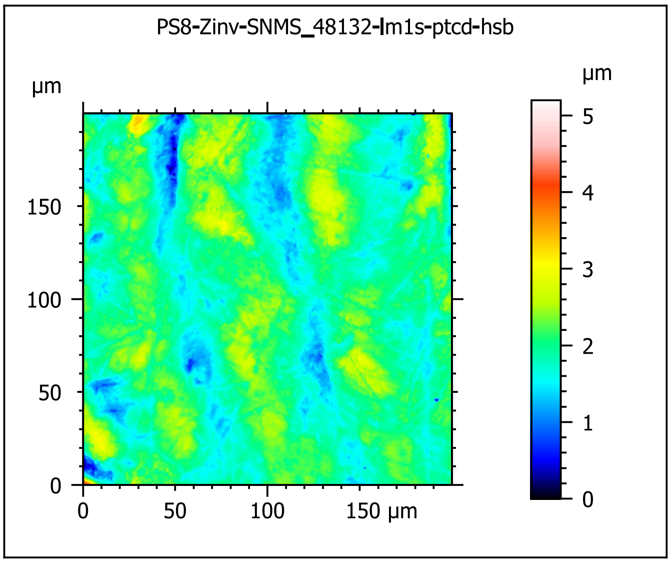


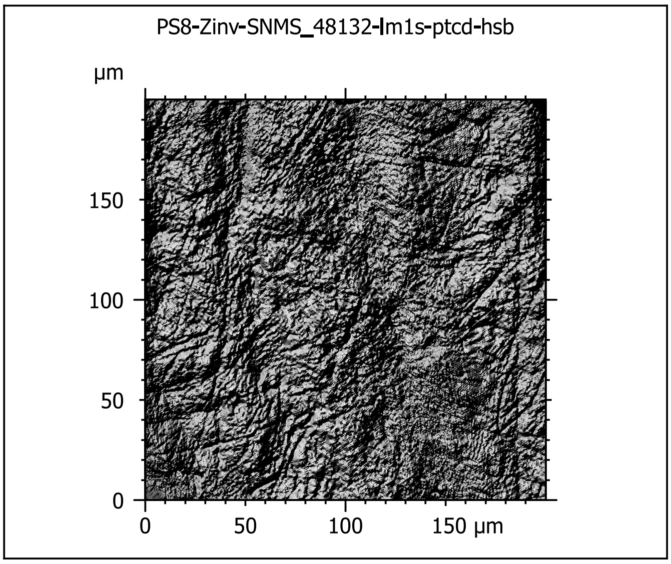


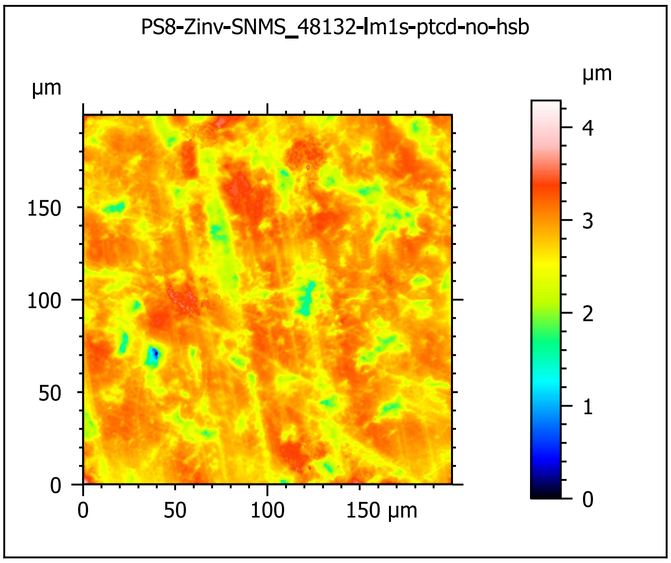


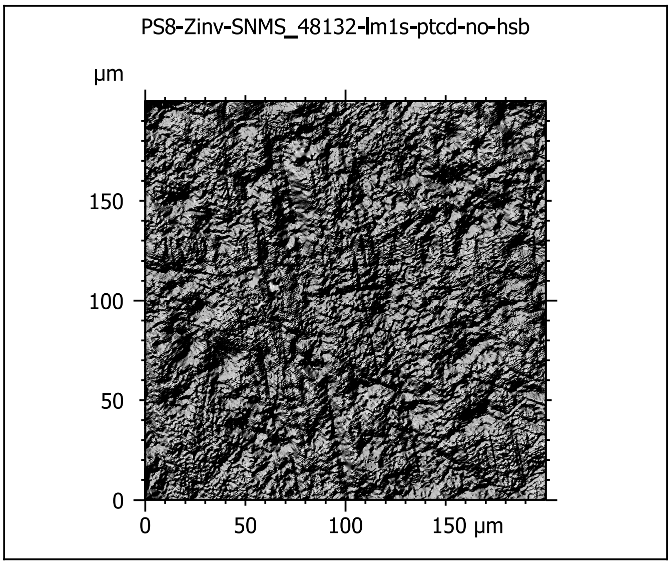


MountainsMap® Premium 10.0.10510

PS8-Zinv-SNMS_48134-lm1s-ptcd-hsb

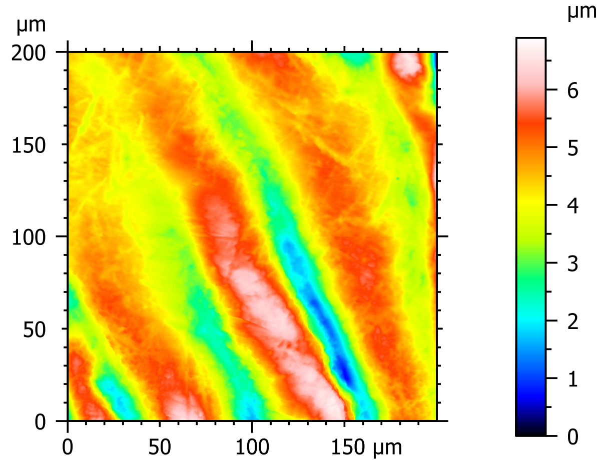


PS8-Zinv-SNMS_48134-lm1s-ptcd-hsb

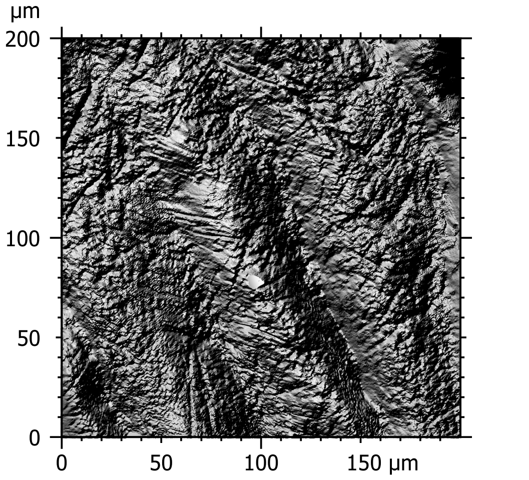


PS8-Zinv-SNMS_48163-UM2s-ptc-hsb

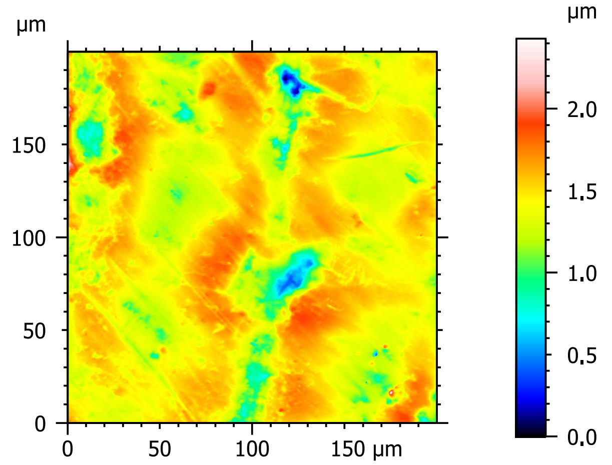


PS8-Zinv-SNMS_48163-UM2s-ptc-hsb

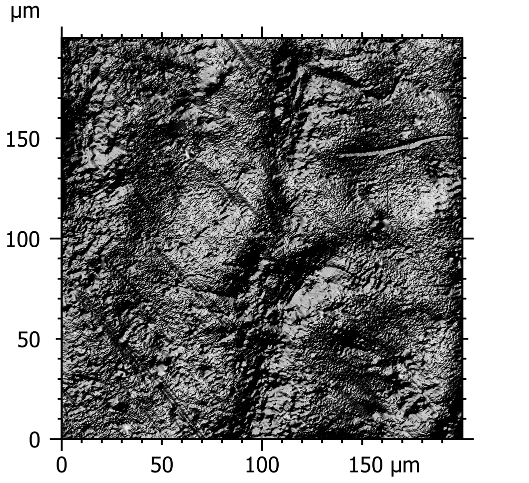


PS8-Zinv-SNMS_48164-UM2d-ptc-no-hsb

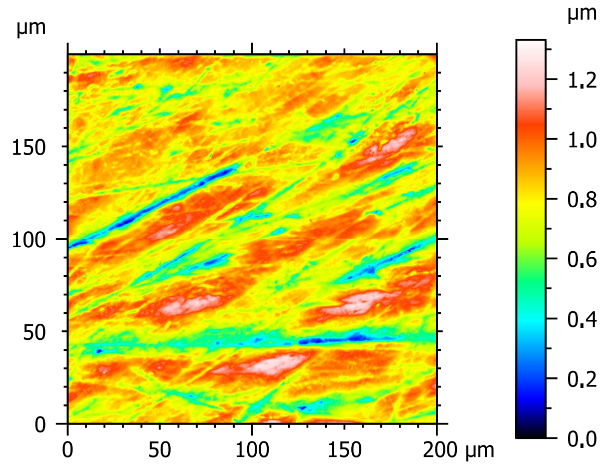


PS8-Zinv-SNMS_48164-UM2d-ptc-no-hsb

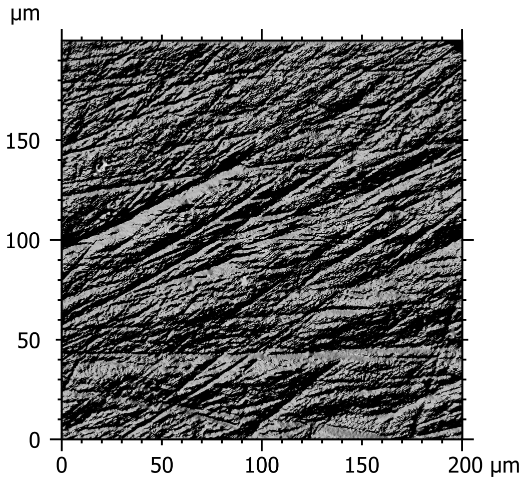


MountainsMap® Premium 10.0.10510

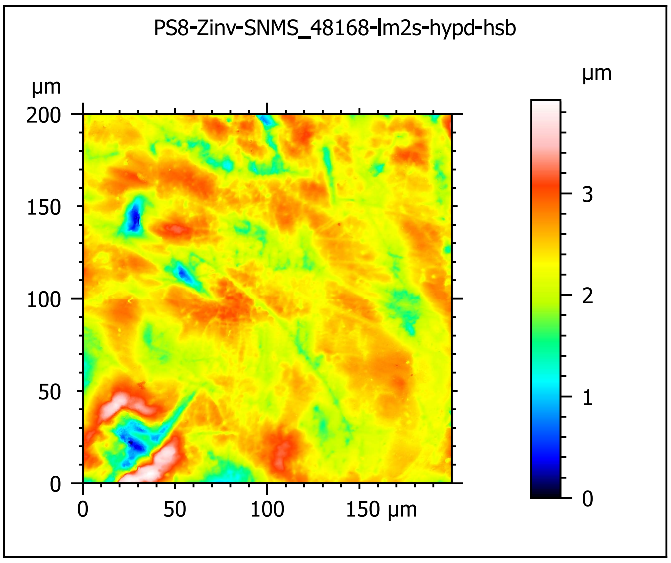


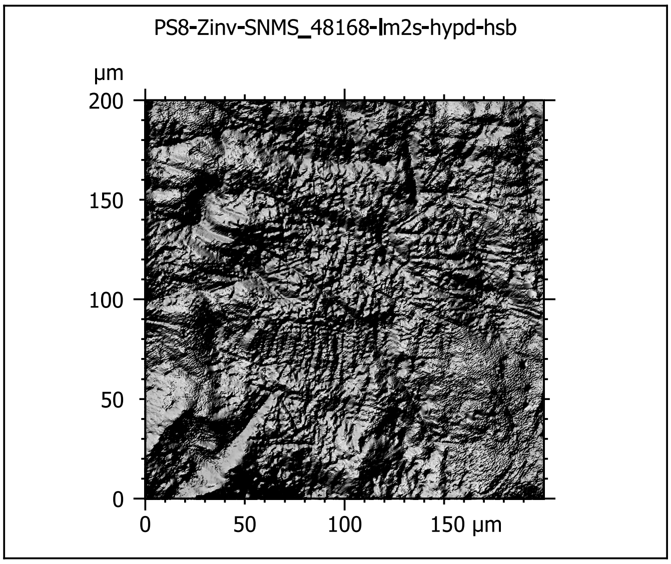


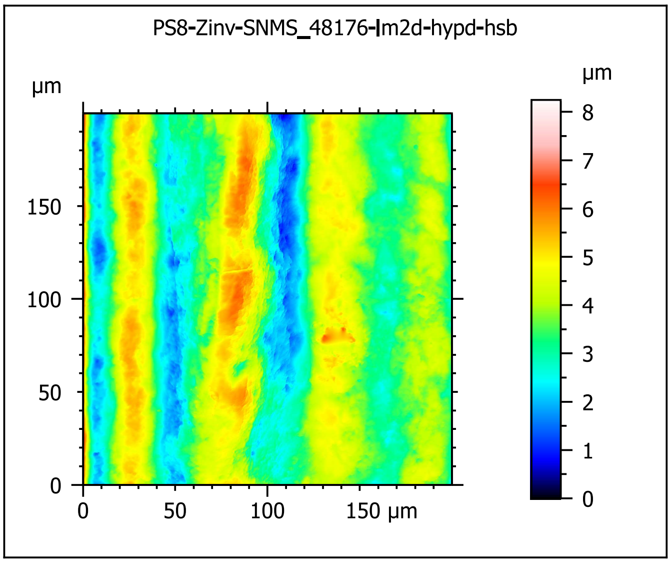


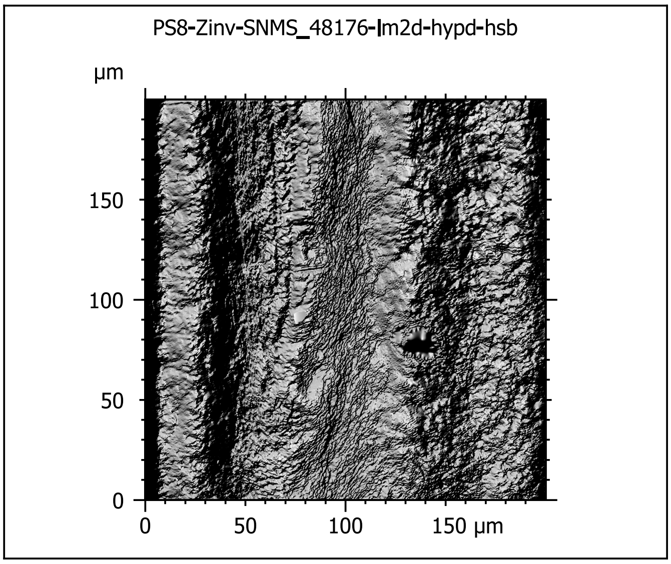


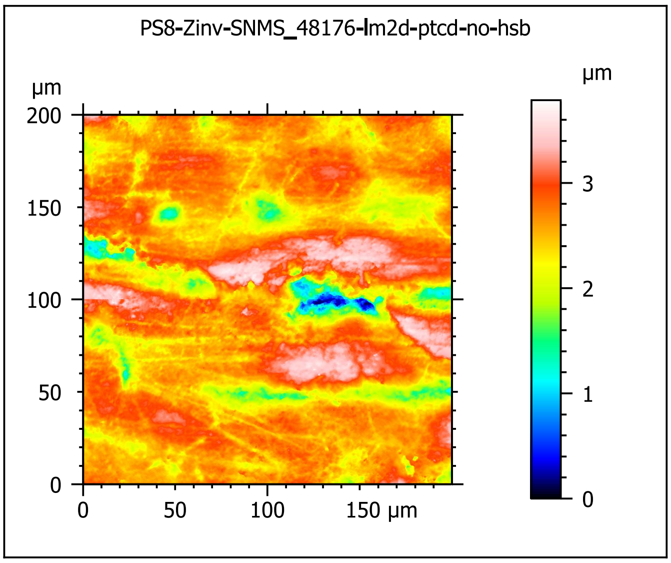


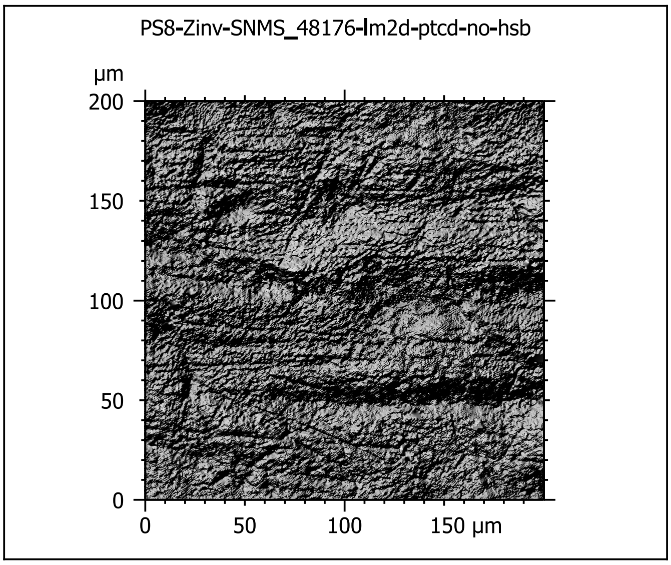


MountainsMap® Premium 10.0.10510
